# Daily rhythms, demographics and health: findings from a nationally representative survey

**DOI:** 10.1101/2023.04.07.536028

**Authors:** Péter P. Ujma, Csenge G. Horváth, Róbert Bódizs

## Abstract

The timing of daily activity in humans have been associated with various demographic and health-related factors, but the possibly complex patterns of confounding and interaction between these has not been systematically explored. We use data from Hungarostudy 2021, a nationally representative survey of 7000 Hungarians to assess the relationship between self-reported chronotype, social jetlag (using the Munich Chronotype Questionnaire), demographic variables and self-reported health and demographic variables, including ethnic and sexual minority membership. Supporting the validity of self-reports, participants with later chronotypes reported the lowest daytime sleepiness at a later clock time. We found that older age, female sex, a more eastward and southward geographical position, residence in a smaller settlement, less education and income, religiousness and cohabiting with small children were associated with an earlier chronotype. Younger age, higher education and income, and cohabiting with small children were associated with increased social jetlag. Of the 48 health-related variables surveyed, the relationship with both chronotype and social jetlag were mostly accounted for by age, sex, and socioeconomic effects, but we identified alcohol consumption, smoking, and physical activity as predictors of both social jetlag and chronotype, while a number of disorders were either positively or negatively associated with chronotype and social jetlag. Our findings from a large, nationally representative sample indicate that both biological and social factors influence chronotype and identified both demographic and health-related variables as risk factors for social jetlag. Our results, however, do not support a causal relationship between light exposure and mental health.

## Introduction

The timing of daily activity (wakefulness) and rest (sleep) is significant in a number of scientific fields, such as ecology (temporal niches; ^1^) differential psychology ^2^, physical and mental health ^3, 4^, as well as somnology/circadian rhythm disorders ^5^. Circadian (∼24 hour) rhythms, present in many species are governed by the master or central clock located in the suprachiasmatic nucleus (SCN, ^6^). The latter coordinates the peripheral clocks which are peculiar to various cell types, tissues, and organs. Humans are a species with clear circadian rhythms ^7^, typically with a peak in activities in daytime hours and a nadir during the night ^8^. However, the precise timing of daily activities varies considerably between individuals for a variety of reasons ^9^. The preferred phase of daily activity of an individual – frequently defined as the individual’s entrainment to the day-light cycle – is defined as the chronotype^10^. Chronotype is associated with both normal and pathological variation in humans.

Younger individuals, especially younger males, are on average characterized by a later chronotype ^8, 11–13^, as are people who live farther to the west within the same time zone and consequently experience sunsets at a later clock time ^11, 12, 14^. Besides evident sex-differences in preferred sleep-wake schedules/chronotypes, sexual orientation was shown to associate with anatomo-functional and phenotypic features of the master circadian clock and timing of activity and rest. Evidence suggests the increased number of arginine vasopressin-containing cells in the suprachiasmatic nucleus (SCN) of homosexual men, paralleled by an increased overall SCN volume ^15^, and data on earlier chronotypes in the members of this sexual minority ^16^. A later chronotype is often associated with worse health outcomes across multiple health domains ^17^, including both mental and physical health. Changes in the timing of daily activities, including the dampening of the daily rhythm, are frequently observed in mental health disorders ^4, 18, 19^. A series of longitudinal studies found associations between chronotype and both mental and physical health outcomes, including mortality ^20–22^ (but see also ^23^).

One of the reasons a late chronotype may be associated with worse health outcomes is that it is may lead to increases in social jetlag, the misalignment between the preferred and socially required timing of daily activity ^24^. Compulsory activities in a modern society, such as school and work typically have an inflexible timing which is more appropriate for individuals with an early chronotype ^25^. Consequently, those who have a late chronotype may experience a large discrepancy between the timing of free day and working day activities, and over the course of many years the repeated need of temporal readjustment may negatively affect health in a broad sense ^26, 27^. In line with this hypothesis, increased social jetlag is associated with worse health outcomes in (see e.g. ^24, 28, 29^ for reviews), including a pseudo-experimental study using time zone boundaries as natural experiments ^30^.

A third factor connecting the timing of daily activities to specifically mental health outcomes is exposure to natural light ^31^. Light therapy is a widely used intervention to treat depression^32^, but some research indicates that more exposure to natural light is associated with better mental health outcomes even in normal populations ^33–35^. However, the relationship between light exposure and mental health may be complex. For example, reduced light exposure may be a consequence rather than a cause for depression if affected people spend more time indoors; or it is possible that low income is associated with both working outdoors (as physical laborers rather than well-paid office employees) and poorer mental health, masking a genuine relationship between light exposure and mental health if other correlates (such as low income) are not held constant.

Former studies on chronotype differences focused on demographic variables ^8^ or health correlates ^3^, but much less on the parallel effects of the above-mentioned sets of variables, leaving the complex picture largely unresolved due to the lack of efficient collection of a substantial data set concerning potential underlying factors. In the current study, we took advantage of a large, nationally representative study to extend and replicate previous literature into these issues. Our goal was to systematically explore what demographic, geographic and health-related human characteristics are associated with chronotype and which of these characteristics are also associated with social jetlag, contributing to the risk of future disease. We also sought to replicate previous literature about the relationship between light exposure and mental health, leveraging a large sample, multiple assessments of mental health, and an analytical strategy which carefully addressed possible confounders.

## Methods

### Hungarostudy 2021

Hungarostudy 2021 was a nationally representative survey launched by the Mária Kopp Institute for Demography and Families and the Institute of Behavioural Sciences at Semmelweis University to assess bio-psycho-social risk factors of disease in the Hungarian population ^36^. It is a follow-up of previous surveys in 1988, 1995, 2002 and 2013 ^37^. In total (not accounting for missingness in specific variables) 7000 participants took part in Hungarostudy 2021. Participants were visited in their households between July 21 and September 15 2021 and filled out questionnaires distribute by an interviewer. The precise date and time of interview completion was recorded. Participants were asked to report their sleepiness level on a Likert scale of 1 to 10 (1 for lowest and 10 for highest sleepiness) before and after interviews ^38^. For analyses estimating the effects of chronotype and administration time on daily sleepiness (see Results), we expressed administration time as hours relative to noon.

After the application of survey weights, Hungarostudy is representative for the Hungarian population in terms of sex, age, education, and place of residence.

### Chronotype assessment

Hungarostudy participants filled out the validated Hungarian version of the Munich Chronotype Questionnaire (MCTQ) to assess their daily patterns and chronotype ^26, 39^. MCTQ is a self-report questionnaire of typical bedtimes and wake times, measured separately on working days and free days. Chronotype is calculated from these reports by taking the sleep midpoint on free days, adjusting for the effects of oversleeping. We use this measure, MSFsc (free-days sleep midpoint adjusted for oversleeping) as the indicator of chronotype. In most our analyses, this value was our dependent variable. We discarded chronotype data from participants who used an alarm clock or indicated having to wake up before their preferred time even on free days. We also excluded participants reporting over 15 hours of daily sleep as likely erroneous entries.

Social jetlag is the difference between the timing of activities on free days and working days^10^. Two types of social jetlag can be calculated from the MCTQ: relative social jetlag, which permits negative values (that is, later timing of activities on working days), and absolute social jetlag, which is the absolute value of relative social jetlag, thus by definition positive.

In analyses seeking the correlates of social jetlag, we used the more expressive relative social jetlag as our dependent variable.

MCTQ also assesses time spent outside on working days and free days. We summed these two values (after multiplying working day exposure by the number of working days and free day exposure by seven minus the number of working days) to obtain a measure of weekly light exposure, which was used as an independent variable. For participants not reporting a regular work-rest schedule and thus a specific number of working days per week, we assumed 5 working days and 2 free days.

### Geographical variables

Participants reported their place of residence at the time of filling out questionnaires. This was cross-referenced with a free web-based database of Hungarian settlements (available at https://webdraft.eu/orszagok_varosok/telepulesek.txt) to fill out for each participant the following variables about their place of residence:

**- Longitude**
**- Latitude**
**- Number of residents.**

Coordinates for the largest settlements were checked for accuracy using https://geohack.toolforge.org/ and some minor discrepancies compared to this database were corrected manually.

### Demographic variables

We considered a set of demographic variables to be used as predictors of chronotype. Participants self-reported their sex as male or female, and their birth year from which their age at interview completion was calculated. Further self-reported variables were the following:

- **education** (fewer than 8 school grades, 8 school grades, vocational schooling, vocation schooling with high school degree, high school degree, university degree, doctoral degree or habilitation). This was re-classified into a three-level variable describing participants’ education level as ‘Basic’ (no high school degree), ‘Intermediate’ (high school degree but no university), and ‘Advanced’ (university or doctorate)
- **sexual orientation** as heterosexual, homosexual or lesbian, bisexual or asexual
- **net monthly personal income**, originally reported in 15 categorical brackets from 0- 30000 HUF to more than 501000 HUF. In order to create a continuous variable from this measure, for the lowest 14 categories we set the value of this variable to the bracket midpoint (for example, for the 0-30000 HUF bracket to 15000 HUF). For the highest bracket which could have included any income level above a threshold, we conservatively set income levels to 550000 HUF.
- **cohabitation**: participants were asked if they are cohabiting with anybody else. Their value on this variable was set to 1 if they reported cohabiting with anybody else and 0 if not. We coded a type of cohabitation, cohabitation with small children, as a separate variable of interest. As Hungarostudy 2021 did not explicitly ask about this, we considered participants to be cohabiting with small children if they 1) reported cohabiting with their children 2) reported having been pregnant in the past six years, acknowledging that this method may misclassify a small minority of participants with adopted children, or their biological children adopted away.
- **ethnicity**: participants self-reported their ethnicity as Hungarian, Gypsy/Roma, German, Romanian, Slovakian, Bulgarian, Greek, Croatian, Polish, Armenian, Rusyn, Serbian, Slovenian or Ukrainian, these categories not being mutually exclusive. 96.9% of participants described themselves as Hungarian and 5.3% as Gypsy/Roma. For the other ethnic categories, we received at most 15 responses, and henceforth excluded them from analyses and only report findings comparing those who reported Gypsy/Roma ethnicity with those who did not.
- **religion**: participants were not asked about their denomination, but they reported the role of religion in their lives on two ordinal items. The first item asked participants if participants practiced any religion (possible responses: not a believer, doesn’t practice religion, practices religion in own way, rarely practices it in church, regularly practices it in church). The second item asked participants how important religion was in participants’ lives (possible responses: not important at all, somewhat important, very important, it influences everything I do). We calculated the sum of these items as a simple measure of religiousness and treated it as a continuous variable.

### Anthropometric and biomedical variables

We considered scores on a series of standardized questionnaires as well as responses to several custom questions as indicators of anthropometric and biomedical characteristics with a potential association with chronotype. Standardized questionnaires were the following:

- **Patient Health Questionnaire-14** ^40^: PHQ-14 is a series of questions assessing 14 common symptoms of pain and problems with the cardiovascular, respiratory, nervous or digestive system. It is identical to the original 15-item version apart from the exclusion of an item concerning sexual problems. Participants can report having each complaint as having “never occurred”, “occurred, but caused no disturbance”, “occurred, and caused some disturbance” or “occurred, and caused major disturbance”. We considered both the sum score on this questionnaire and the responses for each item (dichotomized as 0 for the first two and 1 for the second two answers) as independent variables.
- **Athens Insomnia Scale** ^41^: AIS is a psychometric tool to assess insomnia symptoms. We used five items of the AIS (sleep induction, awakenings during the night, final awakening earlier than desired, sleepiness during the day, well-being during the day) to estimate sleep complaints, and considered the sum score of AIS as an independent variable.
Custom questions were the following:
- **Body Mass Index (BMI):** calculated from self-reported height and weight
- **being treated for one of the following illnesses**: type 1 or 2 diabetes, liver disease, asthma, other respiratory illness, allergy, stomach or intestinal ulcer, other digestive disease, renal disease, rheumatoid arthritis, other disease of the musculoskeletal system, road accident, workplace accident, home accident, high blood pressure, cancer, psychiatric illness, anxiety disorder, heart disease, cerebrovascular disease, COVID-19 or complications, adverse reaction to COVID vaccination, other illness. For each of these illnesses, participants could indicate that they either “did not need treatment”, “their treatment was postponed” (due to the COVID-19 pandemic), “had ambulant treatment”, or “were hospitalized”. We dichotomized all questions to classify participants as not having had an ailment (first response option, coded as 0) or having had it (any other options, coded as 1).
- **alcohol consumption**: participants reported how often they consume alcohol, and (after seeing a short description of alcohol units) how many alcohol units they consume on a typical day when they drink. From this, we estimated monthly alcohol consumption by multiplying frequency (“never” [considered as 0], monthly or more rarely” [considered as 1], “2-4 times per month” [considered as 3], “2-3 times per week” [considered as 10], “four or more times per week” [considered as 16]) with units per drinking occasion.
- **smoking:** we considered participants smokers if they reported smoking cigarettes or electronic smoking devices, and non-smokers if they reported never smoking or having given up smoking.
- **physical activity:** participants reported on two ordinally-scored questions (response options: “never”, “less than once a week”, “once a week”, “many times each week”, “once daily”, “several times each day”) how often they 1) perform at least 30 minutes of exercise, 2) perform at least 10 minutes of other vigorous physical activity (such as gardening or construction). We summed the scores on these items as an estimate of physical activity and treated it as a continuous variable.
- **psychotropic medication use:** participants answered if they are taking any form of psychotropic medication, such as sleeping pills, anxiolytics, or stimulants. Responses could be “never”, “once a month or more rarely”, “4-5 times a month”, “2-3 times a week”, “4-6 times a week”, “once daily”, “several times each day”. We dichotomized the sample as non-users or occasional users (first two response categories) and regular users (all other responses).
- **number of days missed from work due to illness:** as a further estimate of disease burden, participants were asked with a single question how many days they missed from work due to any illness. This response was considered as a continuous variable.

### Well-being and mental health

Hungarostudy 2021 contained a series of standardized questionnaires, the scores on which were considered as predictors of chronotype. These were the following:

- **Abberviated Beck Depression Inventory** ^42^. BDI-9 is a 9-item clinical screening tool for the most common symptoms of depression which is the abbreviated version of the Beck Depression Inventory. We considered the BDI-9 sum score as a continuous independent variable.
- **Perceived Stress Scale** ^43, 44^. PSS is a 10-item questionnaire about the frequency of experiencing stressors and the respondent’s ability to cope with them. We considered the PSS sums score as a continuous independent variable.
- **WHO5** ^45, 46^: The World Health Organization Well-Being Index (WHO-5) is a short self-reported measure of current mental wellbeing.

### Statistical analyses

We investigated the correlates of chronotype and social jetlag in three domains: demographic, geographical and anthropometric/biomedical. Our analytical strategy was slightly different for the three domains, owing to the different logic of interpreting associations. For all models, we used all observations with available data, resulting in different sample sizes across models.

For demographic predictors, we fitted a single linear model with chronotype/social jetlag as the dependent variable and all predictors entered jointly. This is because due to their expected correlations we considered the effects of these predictors most informative when considered net of each other. For example, if men have both later chronotypes and higher incomes, the relationship between chronotype and both sex and income is best interpretable if sex effects are expressed controlling for income and income effects controlling for sex. Because over a thousand participants did not provide income data, we fitted the demographic model both with and without including income as a predictor.

For geographical predictors, we entered all predictors (longitude, latitude and population) jointly, but we ran three models: one without any further controls, one with controls for age, sex and their interactions, and another with an additional control for education and income. This is because while the effects of geographical predictors are the most informative net of each other, it is important if their effects are driven by known demographic effects. Because chronotype was strongly correlated with both relative (r=0.54) and absolute (r=0.52) social jetlag, for social jetlag we added a further model also controlling for chronotype.

For anthropometric/biomedical predictors, we fitted three sets of linear models, each with chronotype as the dependent variable and a single anthropometric/biomedical predictor, either by itself (Model 1), controlling for age, sex and their interactions (Model 2) or additionally also education and income (Model 3). This is because, as in case of geographical predictors, we considered it interesting if associations with anthropometric/biomedical predictors are robust to known demographic correlates of these. In case of social jetlag, as with geographical effects, we added a further model (Model 4) also controlling for chronotype.

When estimating the effect of light exposure on mental health, in addition to unadjusted models, models adjusted for age, sex and their interactions, as well as for education and income, we also adjusted models for settlement type in a fourth model. This is because both mental wellbeing and light exposure can be hypothetically associated with aspects of urban and rural life that are not fully captured by socioeconomic variables.

For chronotype, we express our effect sizes as the expected change in chronotype, in minutes, per unit of change of each predictor. For social jetlag, effect sizes refer to a one our change per unit change of each predictor. For the ease of interpretation, we express BDI, PSS10 and PHQ scores as well as self-reported physical activity and religiosity in z-scores, but the other predictors in natural units.

Code used in our analyses is available in the Supplementary Data.

## Results

### Descriptive statistics

48% of the sample was male and 52% female, with a mean age of 47.9 years (SD=17.3 years). The relative majority reported only basic education (47.8%), 39% reported having intermediate education and 13.2% advanced education. 5.26% of the sample reported Gypsy/Roma ethnicity, at least in addition to Hungarian. For most variables considered in our analyses, over 90% of the 7000 respondents provided valid data, with the exceptions of having been treated for other digestive disorders (valid N=823), income (valid N=4798), chronotype (valid N=4583), social jetlag (valid N=5956) and light exposure (valid N=3606). Frequency tables (for categorical variables) and descriptive statistics (for continuous variables) are provided in the Supplementary data. The Supplementary data also contains detailed results with the exact N used in each model.

### Sleepiness and time of day

Participants were asked about their subjective level of sleepiness (on a scale of 1-10) at both the beginning and the end of interview administration. Because participants could get bored or tired from the interview itself, we used initial scores in the current analyses. In order to validate chronotype measures, we estimated the expected level of sleepiness as a function of chronotype and time of day (relative to noon) at interview administration, including a quadratic term allowing for a parabolic function, with higher sleepiness both in the morning and in the evening. We found evidence for both a linear (-0.048 points per hour, p=5*10**^-^**^4^) and a quadratic (0.029 points per hour squared, p=2*10**^-^**^7^) effect of time of day on self-reported sleepiness, indicating that sleepiness is mainly a parabolic function of time of day. Chronotype was not significantly associated with increased sleepiness (0.001 points per hour, p=0.08). Crucially, the chronotype*squared daytime interaction was significant (p=6*10**^-^**^4^), indicating that the parabolic course of sleepiness over the day is affected by chronotype. Individuals with earlier than average chronotypes reached the lowest level of sleepiness before noon, while those with than average chronotypes during the afternoon (**Figure 1**).

**Figure 1.**
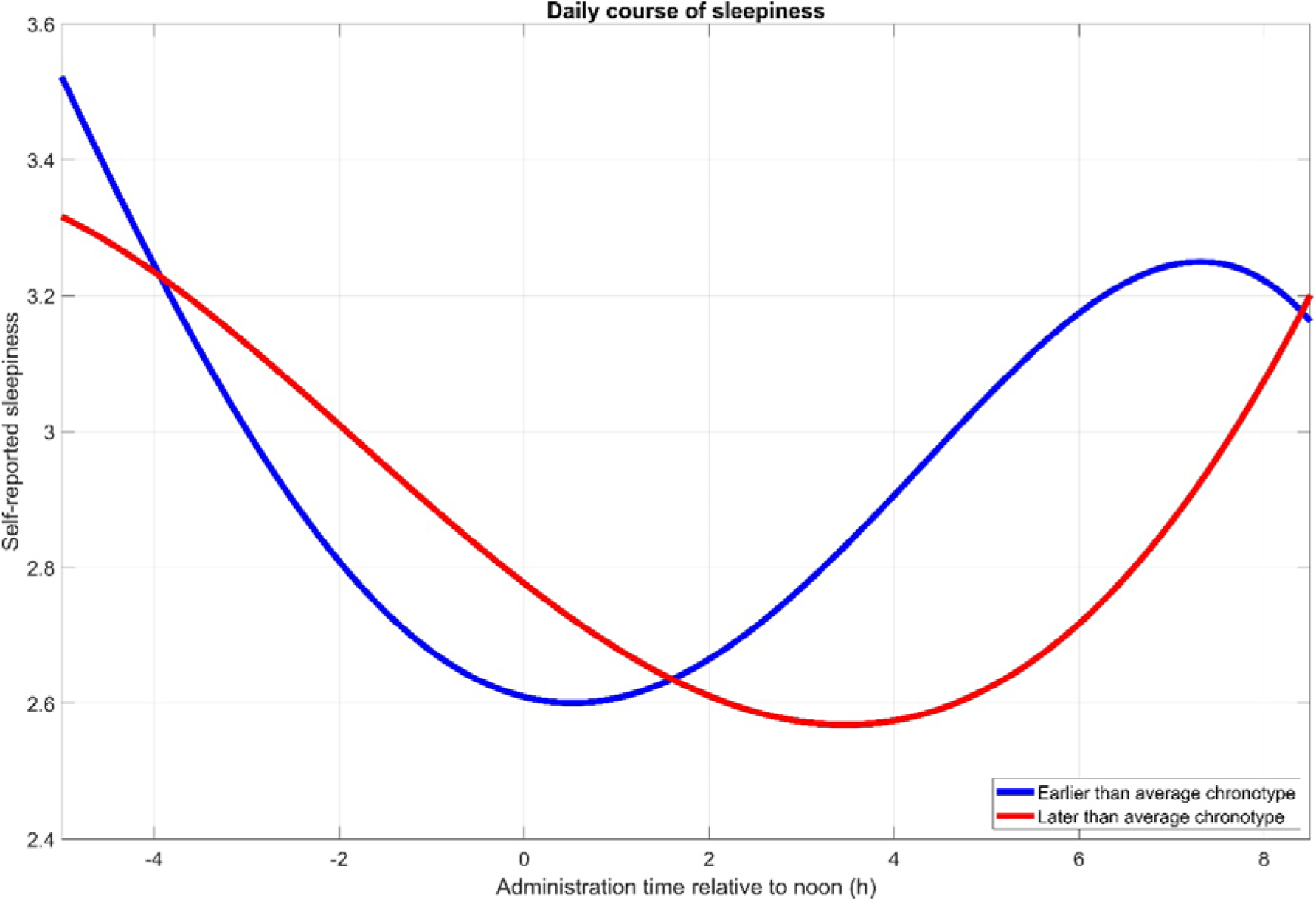
Fourth-order polynomial fit lines of self-reported sleepiness at the beginning of interview administration as a function of clock time, shown separately for participants with earlier or later than average chronotypes. Note the different timing of nadirs. Note that axis X has been truncated at [-5 8.5] (administration time before 7 AM or after 8:30 PM). Fit to participants with such extreme values (N=63) is not shown as it is based on little information, however, the accuracy of these administration times was confirmed, and this data was also used in modelling.

### Demographic associations of chronotype

The reported chronotype was 3.03 hours, corresponding to midsleep just after 3 AM. Several demographic factors exhibited independent associations with chronotype (all p<0.001 unless otherwise specified). Females (by 23.6 minutes) and older participants (by 1.8 minutes per year) reported earlier chronotypes, with a significant sex*age interaction (B=0.3, p=0.016). That is, older age was more strongly associated with an earlier chronotype in men (corresponding to a 0.3-minute larger drop in each year), and the sex difference in chronotype tended to diminish in older age.

Better educated participants also had later chronotypes, corresponding to an 18.6 minute increase from basic to intermediate education and a further 3.5 minutes to advanced education.

Cohabiting with another person by itself had no association with chronotype. However, cohabiting with small children (<6 years old) was one of the strongest predictors of earlier chronotypes, corresponding to a difference of 29.6 minutes.

We found no evidence that ethnic minority members had different chronotypes. Of the sexual minorities surveyed, bisexuals had a notably later chronotype (by 50.6 minutes), with no difference between heterosexuals, homosexuals and asexuals.

More religious people had earlier chronotypes (by 8 minutes per standard deviation).

When added as a predictor, income had an independent positive association with chronotype, with approximately 4.4 minutes later chronotype per 100k HUF of monthly personal income. In these models all other predictors retained their significance with similar regression coefficients. Controlling income, Gypsy/Roma ethnicity was also associated with later chronotype (15.3 minutes, p=0.024).

Findings about demographics effects are summarized in **Figure 1**. Detailed model results are reported in the Supplementary data.

### Geographical associations of chronotype

Hungarostudy 2021 sampled individuals across the entire area of Hungary. In line with previous findings, we found that latitude, longitude and the size of the place of residence had independent associations with chronotype (all p<0.001), with individuals living farther to the west (8.7 minutes per degree), farther to the north (9.3 minutes per degree) and in larger settlements (1.6 minutes per 100k inhabitants) having later chronotypes (**Figure 2**). These effects were robust to controlling for age, sex, education and income. Detailed statistics are reported in the Supplementary data.

**Figure 2.**
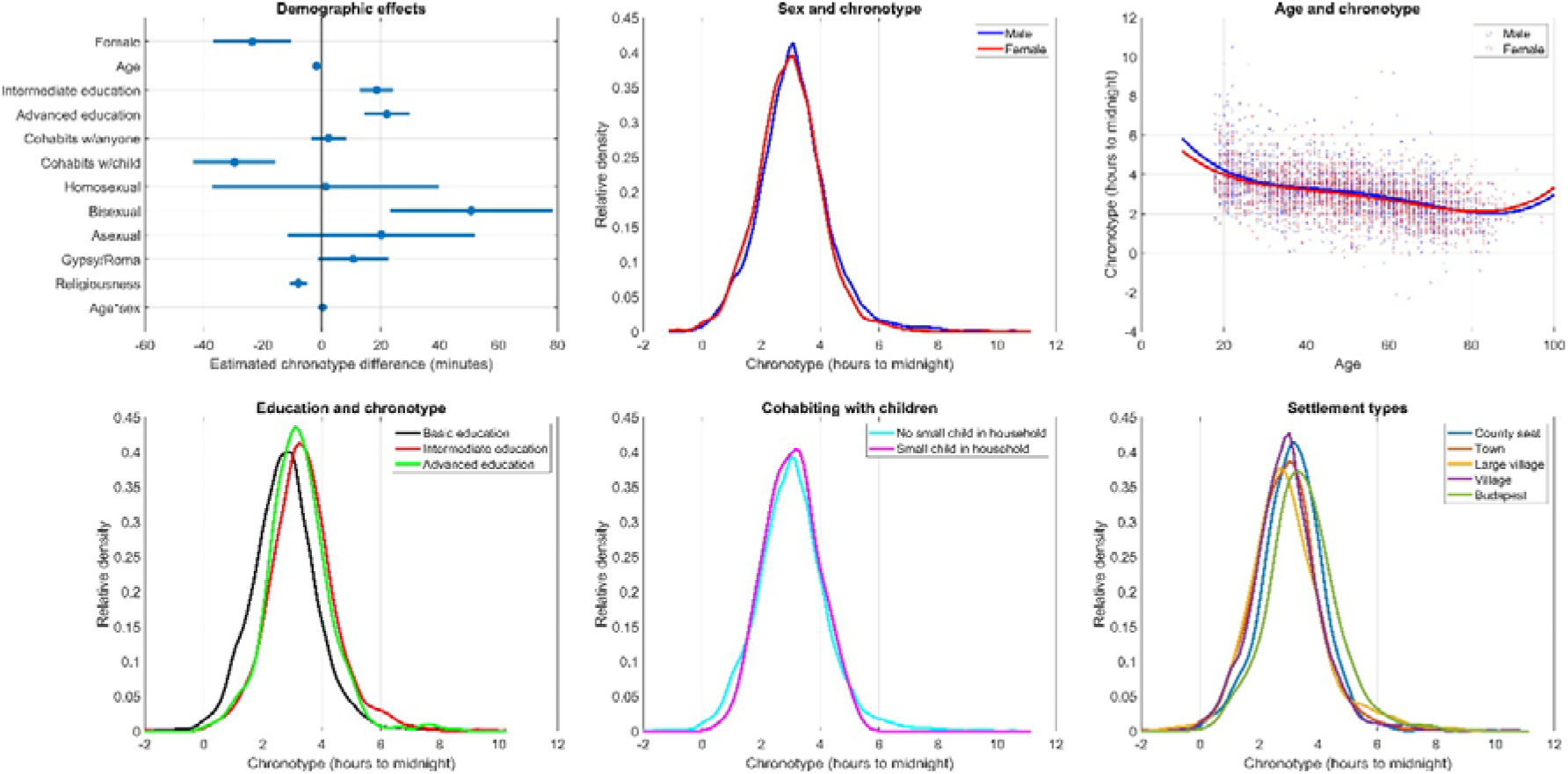
Demographics effects on chronotype. **Panel A**: partial regression coefficients with 95% confidence intervals. Regression coefficients indicate the expected change in chronotype in minutes as a function of one unit change in the predictor (for continuous variables) or relative to the reference category not shown (for categorical variables), net of the other predictors. **Panel B**: Kernel density estimates of chronotype in males and females. **Panel C**: The diminishing sex difference in chronotype at older ages. Data points are overlain with 4**^th^** order polynomial fit lines. **Panel D**: Kernel density estimates of chronotype in the three educational categories. **Panel E**: Kernel density estimates of chronotype as a function of cohabiting with small children. **Panel F**: Kernel density of chronotype as a function of place of residence. “Large village” corresponds to a legal category (nagyközség) in the Hungarian administrative system.

We calculated a settlement-level chronotype by averaging the chronotypes of all participants providing data from each settlement (N=304). In settlement size-weighted linear regression, we used this settlement-level chronotype as the dependent and longitude, latitude and settlement size as independent variables. We found that geographical effects on chronotype replicate on the settlement level. Hungarian settlements further in the west of the country (4.8 minutes/degree, p=1.6*10**^-^**^5^, **Figure 3**) and with larger populations (1.32 minutes/100k inhabitants, p=8*10**^-^**^12^) had later average chronotypes. While the size and direction of the effect size was similar to participant-level analyses, we found no significant effect for latitude (B=3 minutes/degree, p=0.28), possibly due to lower power.

**Figure 3.**
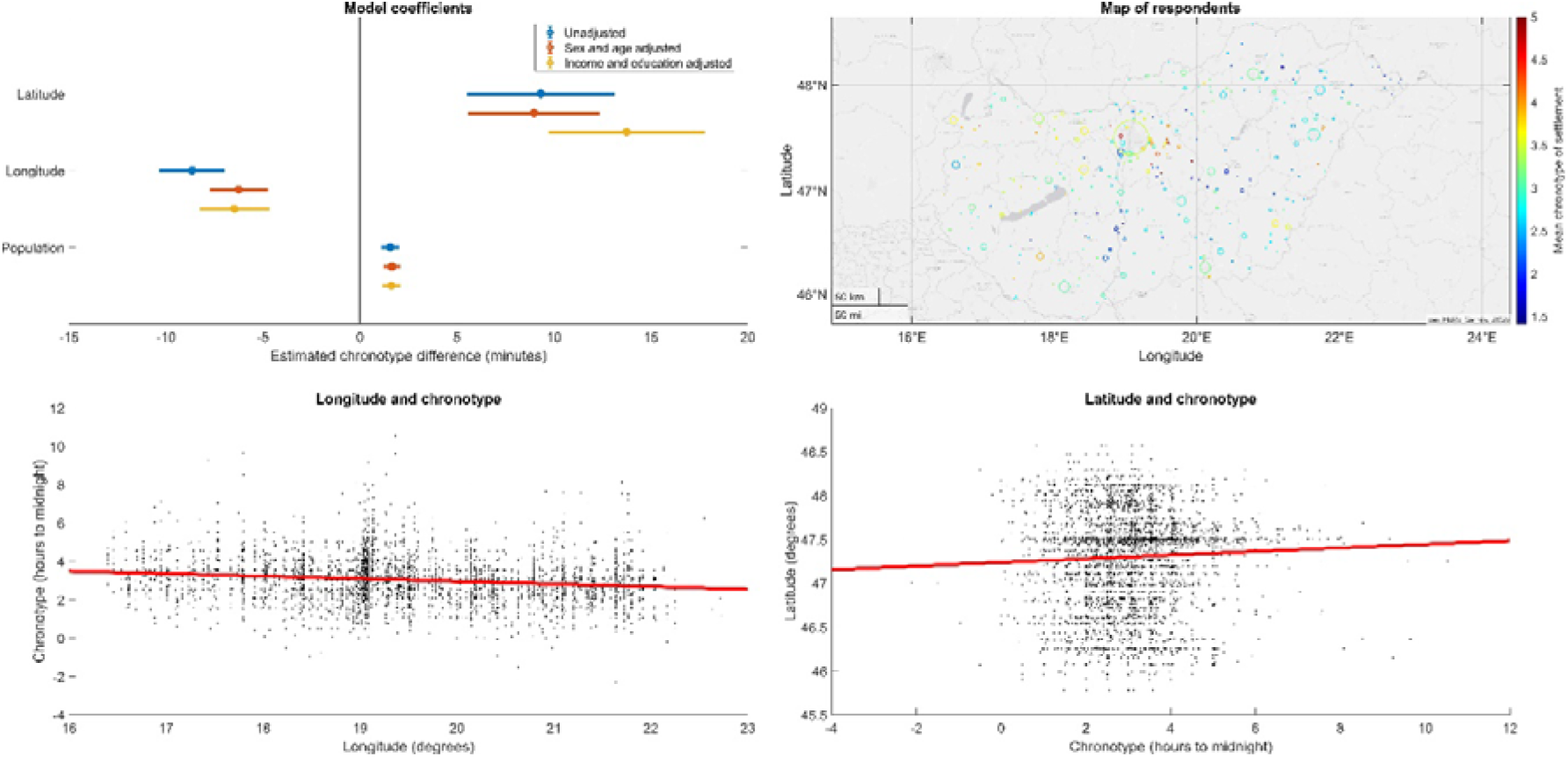
Geographical effects on chronotype. **Panel A**: partial regression coefficients with 95% confidence intervals. Regression coefficients indicate the expected change in chronotype in minutes as a function of a one-degree change in longitude/latitude or per 100k inhabitants. **Panel B**: the distribution of participants over the territory of Hungary. The size of the circles is proportional to the number of responses received from each settlement. Color coding indicates the mean chronotype of settlement. **Panel C**: the association between longitude (west-east) and chronotype. **Panel D**: the association between chronotype and latitude (south-north). Axes on Panel C and D are oriented so that geographical position is shown as it usually appears on maps.

A visual inspection of **Figure 4** suggested that the relationship between settlement population and chronotype is different as a function of longitude (note that in more eastern longitudes larger settlements tend to increasingly fall above the regression line). Therefore, we tested another model adding a longitude*population interaction. This interaction effect was indeed positive but not significant at the settlement level (p=0.19). However, it was highly significant on the individual level with greater power (B=5.4, p=2*10**^-^**^6^), suggesting that in Eastern Hungary the trend of respondents in larger settlements having later chronotypes is amplified. This may reflect that the urban-rural divide in lifestyles is larger in this less economically developed part of the country.

**Figure 4.**
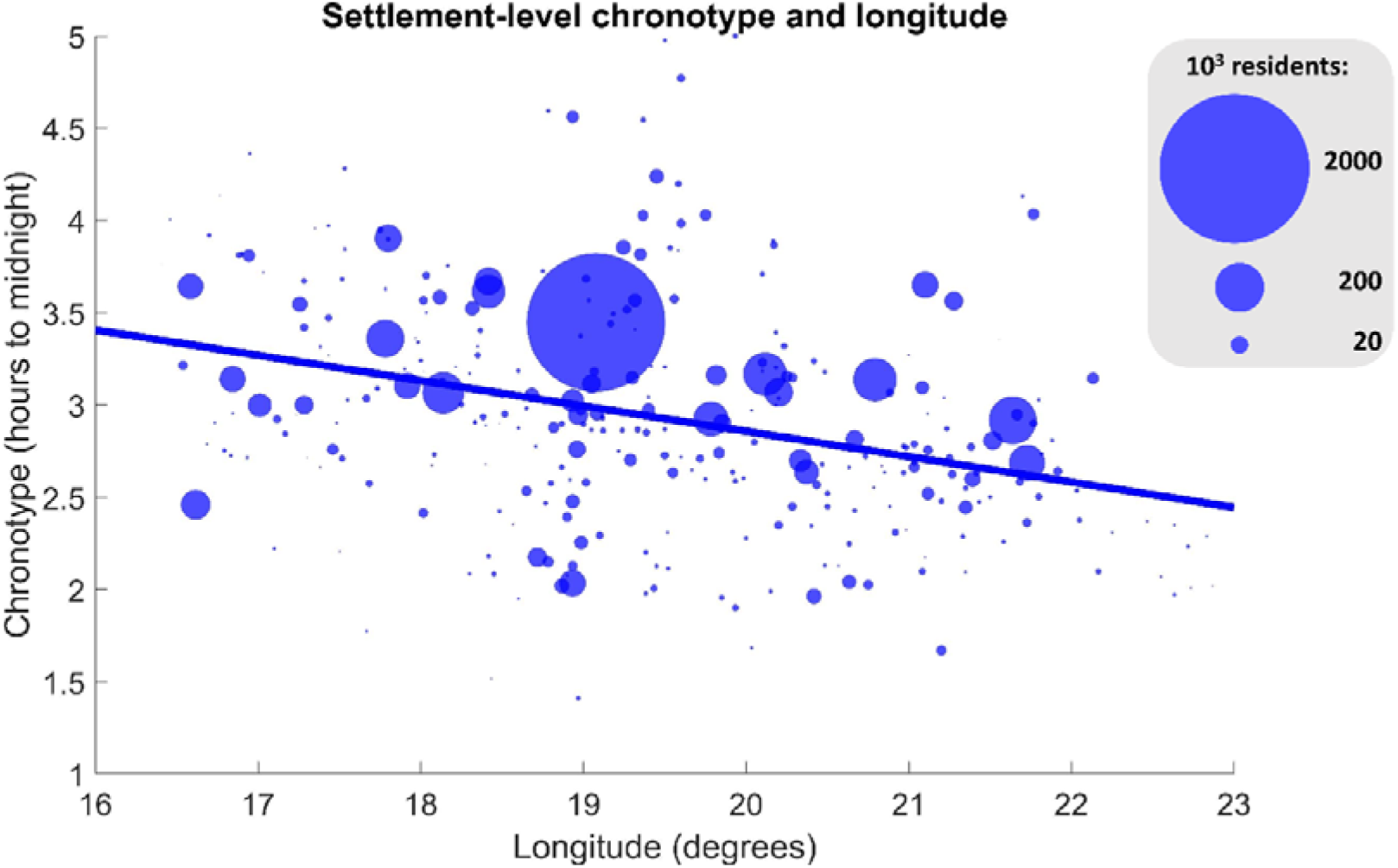
The correlation between settlement-level average chronotype and longitude. Data points show the mean of all chronotype estimates from each unique settlement surveyed in relation to their geographical position. The size of data points is proportional to settlement size. The unweighted least-squares fit line is shown.

**Figure 5.**
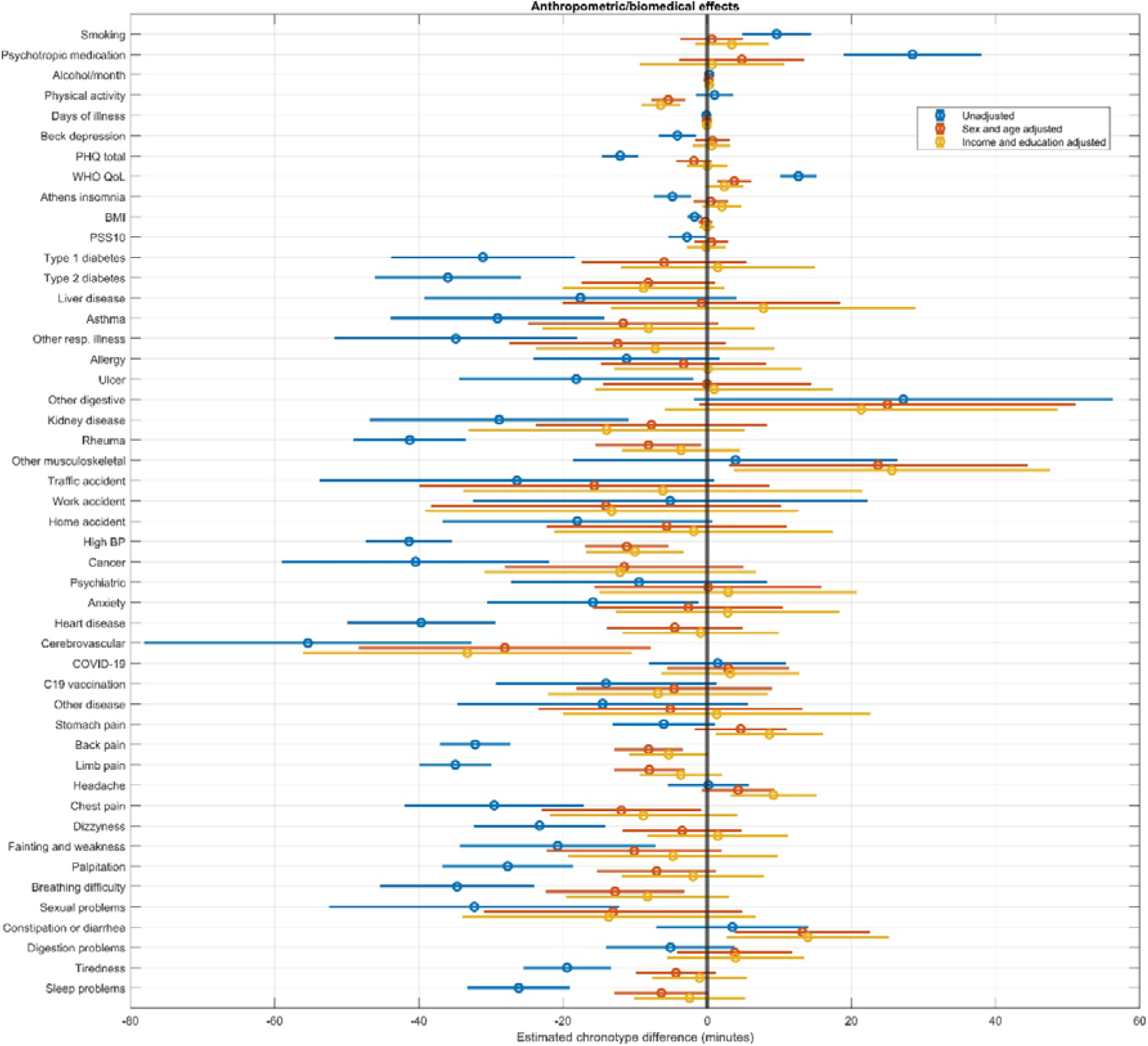
Associations between anthropometric-biomedical indicators and chronotype. The figure shows regression coefficients and 95% confidence intervals, either for univariate models (blue) or after adjustment for sex and age (red), or additionally also for income and education (yellow). Regression coefficients indicate the expected change in chronotype in minutes as a function of a one unit change in the predictor (for BMI, alcohol consumption and mental health questionnaire scores) or in those suffering from an ailment relative to those who do not (for all others).

**Figure 6.**
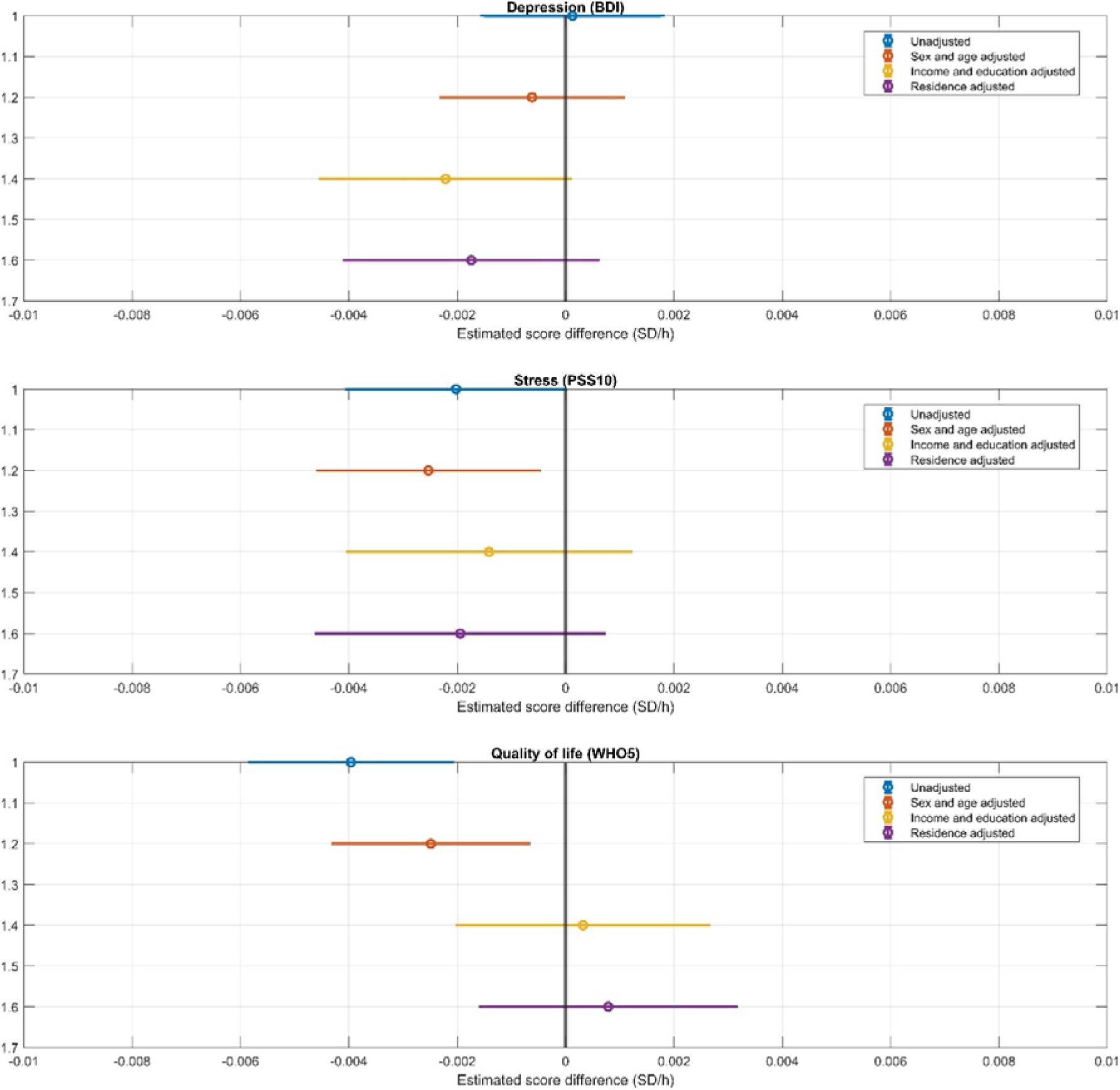
The association between light exposure and three measures of mental health: scores on the Beck Depression Inventory (top), the Perceived Stress Scale (middle) and the WHO Well-Being Scale (bottom). Markers indicate regression coefficients (expected score change in standard deviation units per an hour change in weekly light exposure) and 95% confidence intervals. Four models, with increasingly strict covariate sets are illustrated.

### Anthropometric and biomedical associations

Next, we investigated if chronotype is associated with a set of anthropometric, biological and medical predictors. In univariate models, most anthropometric and biomedical variables were associated with chronotype, most typically with an earlier chronotype. Notably, even in these models, having been treated for psychiatric problems, COVID-19 or complications of COVID-19 vaccination were not associated with chronotype.

Most associations were, however, eliminated after controlling for age and sex, which are strongly associated with both chronotype and disease propensity. Models further adjusted for education and income yielded similar results. Even after adjusting for age, sex, education and income, alcohol consumption, WHO wellbeing score (p=0.029), other musculoskeletal disorders (p=0.019), stomach pain (p=0.014), back pain (p=0.035), headaches and constipation/diarrhea (p=0.011) were associated with a later chronotype, while physical activity, high blood pressure and cerebrovascular disease were associated with an earlier chronotype. (P-values exceeding 0.01 are highlighted as possibly spurious.)

Importantly, we found no associations between chronotype and BMI, the self-reported use of psychoactive medications (sleeping pills, anxiolytics and stimulants), COVID-19 or vaccination complications, smoking or days missed from work due to an illness.

### Light exposure and mental wellbeing

In our next analyses, we assessed whether self-reported weekly light exposure is related to mental wellbeing, using three different self-report measures of the latter and four different covariate sets. In line with previous research ^26^ (but see also ^47^), more light exposure was significantly related to an earlier chronotype (r=-0.179, p=6*10**^-^**^22^), highlighting the validity of this self-reported measure. Scores on the Beck Depression inventory were not significantly associated with light exposure regardless of which (or whether any) covariates were used. Lower perceived stress was associated with light exposure in unadjusted models (0.002 SD lower scores per each hour of weekly light exposure, p=0.045) and if age and sex were controlled for (-0.0025 SD/h, p=0.013), but not after adjustment for income, education and place of residence. Well-being, assessed by WHO5, was associated with a lower level of weekly light exposure (-0.004 SD/h, p=2*10**^-^**^5^). This association weakened after adjusting for adjusting for age and sex (-0.0025 SD/h, p=0.006), but was rendered insignificant after further adjusting for income, education, and place of residence.

Results are summarized on **Figure 4** while detailed statistics are provided in the Supplementary data.

### Correlates of social jetlag

We found that similar demographic factors are associated with social jetlag as with chronotype. Age (-0.025 hours per year, p=10**^-^**^63^) and female sex (-0.39 hours, p=2*10**^-^**^4^) were associated with less social jetlag, with a significant age*sex interaction (B=0.004, p=0.013) suggesting a diminishing sex difference with age. Cohabiting with a small child (-0.49 hours, p=10**^-^**^8^) and religiousness (-0.07 hours per standard deviation, p=10**^-^**^4^) were also associated with less social jetlag. Asexuals (-0.5 hours, p=0.038) and Gypsies/Roma (-0.19 hours, p=0.034) reported less social jetlag. Income, when entered in the model, was related to higher social jetlag (0.17 hours/100k HUF, p=10**^-^**^12^), also introducing a significant effect of advanced (but not intermediate) education, which was related to less social jetlag net of income (-0.19 hours, p=0.01), but reducing minority effects to insignificance.

Participants residing in more westward longitudes (-0.1 hours per degree, p=2*10**^-^**^15^) and more northern latitudes (0.15 hours per degree, p=7*10**^-^**^7^) reported higher levels of social jetlag. Population at the place of residence, however, was unrelated to social jetlag. These associations were not affected by controls for age, sex, education, and income.

We found that several anthropometric and biomedical variables were related to social jetlag. Somewhat surprisingly, several diseases were related to lower levels of social jetlag, and two positive factors (higher self-rated quality of life and lower perceived stress) were related to higher levels. Once age, sex, education, and income were controlled for, 5 (smoking, alcohol consumption, WHO5 life satisfaction, headaches and constipation/diarrhea [p=0.012]) anthropometric and biomedical variables were related to higher levels of social jetlag, while 10 (physical activity, depression, perceived stress, type 2 diabetes [p=0.44], asthma, allergy, kidney disease [p=0.036], traffic accidents [p=0.031], cerebrovascular disease and sexual problems [p=0.03]) were significantly related to less social jetlag (**Figure 7**). (P-values exceeding 0.01 are highlighted as possibly spurious.)

**Figure 7.**
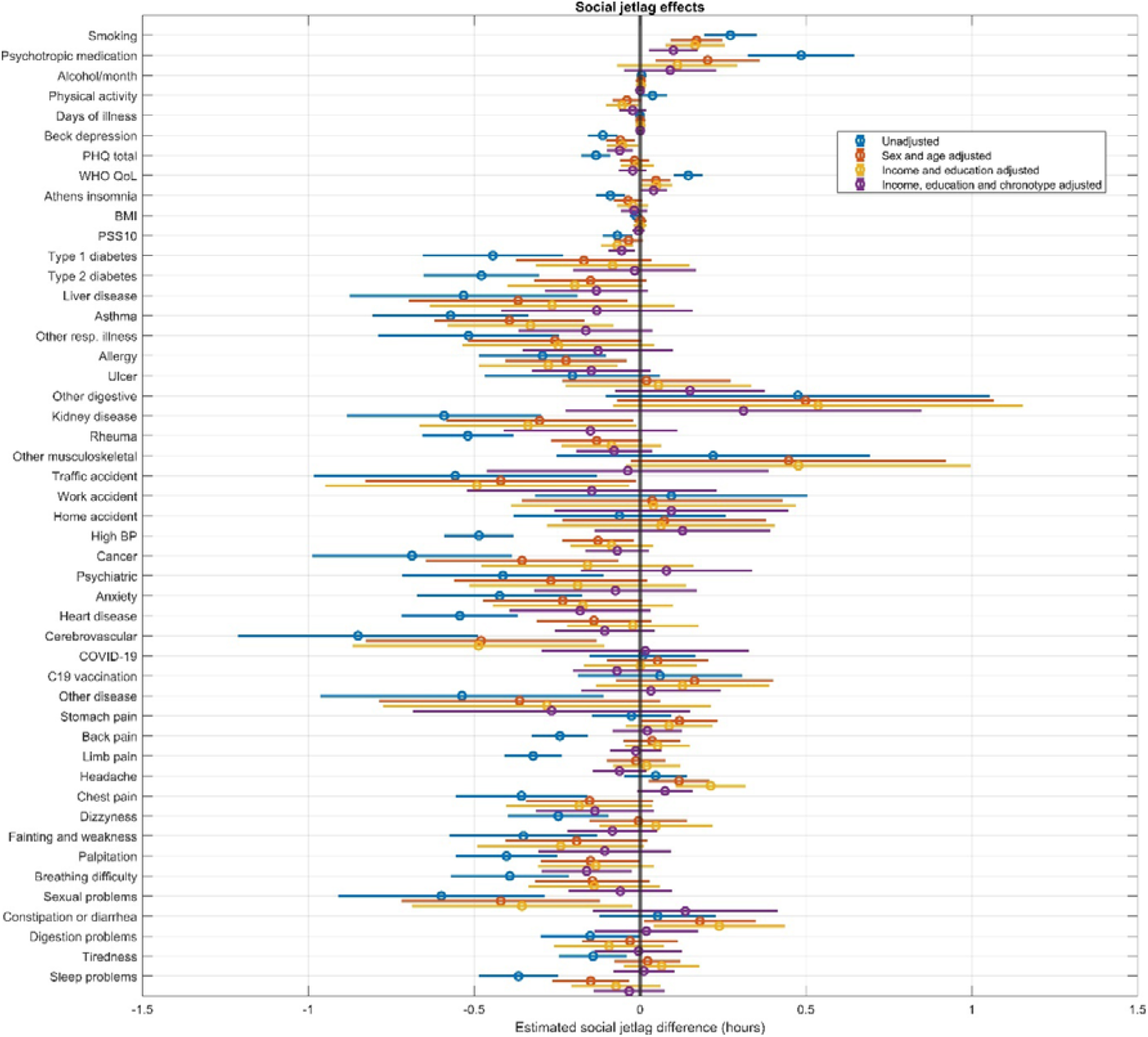
The cross-sectional association between social jetlag and anthropometric/biomedical factors. The figure shows regression coefficients and 95% confidence intervals, either for univariate models (blue) or after adjustment for sex and age (red), for income and education (yellow) or additionally also for chronotype (pink). Regression coefficients indicate the expected change in social jetlag in hours as a function of a one unit change in the predictor (for BMI, alcohol consumption and mental health questionnaire scores) or in those suffering from an ailment relative to those who do not (for all others).

Increased social jetlag was highly (r=0.54 for relative and r=0.52 for absolute SJL) correlated with a later chronotype, suggesting that the correlates of social jetlag may simply reflect differences in chronotype. Controlling for chronotype, age and cohabiting with children was still associated with less social jetlag and income with increased social jetlag. However, the effect of education was reversed, with both intermediate and advanced education being associated with less social jetlag. (Note that chronotype-corrected calculations only include participants with valid chronotype estimates, for example, only people with small children who still report being able to freely choose their weekend wake-up times.) Of geographical effects, only lower longitude (p=0.018) was slightly associated with SJL net of chronotype. Controlling for chronotype completely changed the associations between health and SJL. More social jetlag was associated with smoking and better self-rated mental health (all three questionnaires), with borderline significant associations with headaches (p=0.043) and the absence of palpitations (p=0.018), but none of the other anthropometric/biomedical measures.

Detailed statistics about the correlates of social jetlag are reported in the Supplementary data.

## Discussion

Chronotype refers to the phase of entrainment between a person’s daily sleep/wake cycle to solar time^10^ (the latter roughly approximated by clock time, and this entrainment often measured by self-report questionnaires). Individual differences in chronotype also suggest differences in the timing of optimal vigilance, work capacity and other variables which may interact with demographic factors, social requirements, and health. Although many studies analyzed the demographic, behavioral and health-related associations of chronotype, these potentially overlapping and interacting effects were never analyzed in unison. In this study, we leverage a recent nationally representative survey to provide an exhaustive analysis of the demographic, psychosocial and health-related factors associated with chronotype, exploring how key covariates may alter these associations.

### Chronotype, time of day and sleepiness

Findings from our population-based, highly ecologically valid study cohere with earlier laboratory investigations based on repeated measurements on low sample sizes, but well-controlled settings ^48^ or the outcomes of modelling studies ^49^. Data indicates a U-shaped course of sleepiness during the day, with the lowest population-level average subjective sleepiness in the middle of the day (around noon), whereas higher sleepiness ratings emerge at earlier and later testing sessions. These are hypothesized to reflect sleep inertia and increased sleep pressure, respectively ^49^. Moreover, our results indicate that subjects with an earlier chronotype are characterized by earlier nadir in sleepiness as compared to participants with later chronotypes. This latter finding fits well with the concept of chronotype, as well as with the definition of morningness-eveningness as preferred time schedules or the degree to which respondents are active and alert at certain times of day^50, 51^.The finding that sleepiness is lowest during the midday, but the timing of its nadir is affected by an individual’s chronotype, may be significant in the prediction and prevention of sleepiness-related accidents.

### Chronotype and demographics

In our first analyses we investigated the factors that underlie individual differences in the timing of daily activity. In line with previous studies ^8, 13, 52^, we found that demographic factors are key predictors of chronotype. We found that sleep timing is earlier in older individuals, and while women on average have earlier chronotypes, this sex difference becomes smaller with age. Income and education had largely independent associations with a later chronotype. We found no evidence that individuals living alone have different chronotypes than those who cohabit with others in the same household, however, those with small children reported substantially earlier chronotypes.

To our knowledge, our study is the first large, nationally representative study to specifically investigate chronotype in sexual and ethnic minority populations. Net of other demographic predictors, we found no evidence for substantial differences in sleep timing in the largest ethnic group in Hungary, Gypsy/Roma people and only limited differences in members of the surveyed sexual minorities, specifically a later chronotype in bisexuals, possibly reflecting previously documented individual differences in sensation seeking ^53^. The lack of a replication of earlier chronotypes in homosexuals ^16^ might reflect biased sampling of former non-population-based studies, changes in the biological and socialization-related factors contributing to homosexuality, or simply low power as we only had 18 homosexual respondents in our sample. Limitations of sample size also prevented us from identifying possible sex differences in the chronobiological correlates of homosexuality. We found earlier chronotypes in those reporting higher levels of religiosity. Similar findings were reported in former studies conducted on various populations ^54^, suggesting the cultural invariance of this association.

Overall, these findings indicate that biological and social factors interact to determining sleep timing. While the effects of age and sex as well as their interactions likely reflect the prevailing effects of underlying biological factors, other effects indicate that the different lifestyles of more educated and more affluent respondents include later sleep timing. Childcare is one of the strongest factors associated with an earlier chronotype, most likely to accommodate the needs of children who themselves have a chronotype earlier than adults ^55^.

### Chronotype and geography

In line with previous studies ^11, 12, 14, 56^ we confirmed the observation that even within the same time zone with identical social time, people living further to the east and consequently experiencing earlier sunrises and sunsets report earlier chronotypes. The magnitude of this effect was 8.7 minutes per degree (111 km), which, extrapolated over the west-east span of Hungary of over 500 km, matches or even exceeds the maximal within-country difference in sunrise and sunset times, which is approximately 25 minutes. Data from other countries suggests that the entrainment of human daily rhythms to sun time may surpass country borders. While all Central Europe has the time zone, studies from countries much farther to the west reported chronotypes about as much later as differences in solar time relative to Hungary would suggest, or possibly even more. Compared to our estimate of a mean chronotype of 3.03 hours, for example, a Czech study reported a mean chronotype of 3.1 hours after midnight ^8, 11^, and a large study of German, Swiss, Dutch and Austrian respondents ^8^ reported a mean chronotype at 4:14 AM. Although the distribution of respondents within each country affects the exact mean solar time they are exposed to, a rough evaluation of the figures above can be made by comparing the solar times of country capitals. Budapest has a sunrise 24 minutes before Prague, 35 minutes before Berlin, 54 minutes before Zürich and 68 minutes before Amsterdam, and the magnitude of these differences is similar to chronotype differences reported between respondents in the corresponding countries. (Calculations of solar time are from www.timeanddate.com.) Further studies considering the local times of average work start (e.g. ^57^) are warranted, and policymakers should be wary that solar times varies widely across the European Union, affecting the daily rhythm of people even the clock time is identical.

In line with previous Czech ^11^ and a multinational ^47^ study, but not with a global investigation ^12^, we found that residence at northern latitudes is also associated with a later chronotype. While we cannot account for this unexpected finding, is not confounded by age, sex, education, income or population size, which were controlled for in statistical models. Further investigations are needed to replicate and explain this finding.

### Chronotype and health

In apparent contrast with available reports in the literature revealing health hazards of later chronotypes or eveningness ^17, 58^ most health problems in our current nationwide representative study were associated with an earlier chronotype. This finding was severely confounded by age, as older participants tended to report both more illnesses and earlier chronotypes. Several associations survived correction for age, sex, education and income, however. After these corrections, most health problems (alcohol consumption, stomach pain, headaches, constipation/diarrhea, other musculoskeletal disorders) were associated with a later chronotype which partially coheres with the literature. These health conditions interfere with sleep initiation, while alcohol consumption may be associated with a lifestyle characterized by later bedtimes. Interestingly, not only physical activity as a health-promoting factor, but also two conditions of the cardiovascular systems (high blood pressure and cerebrovascular disease) were associated with an earlier chronotype. While some studies exist that investigated chronotype as a risk factor for cardiovascular disease ^59, 60^, all studies revealed eveningness or later chronotypes to be associated with increased hazards. Indeed, morning chronotypes were associated with increased blood pressure in subjects with apnea– hypopnea index ≥ 15 events per hour ^61^. To our knowledge, neither of the reports had a design comparable to ours. Therefore, our findings require replication.

### No evidence that light exposure improves mental health

Exposure to bright light is a well-established therapy for seasonal depression ^32^ and some previous research has found ^31^ that mental wellbeing is generally associated with light exposure. We investigated this pattern by considering the possible patterns of confounding in a cross-sectional observational sample such as Hungarostudy 2021. For example, it is possible that while light exposure does lead to improved mental wellbeing, most natural variation in light exposure comes from work-related sources such as commuting on foot or working in agriculture, construction or delivery services. Respondents with such characteristics may possess lower average levels of education, income and more frequently live in smaller settlements, which in turn are related to lower mental wellbeing, causing an overall zero association between light exposure and mental wellbeing in an observational study. Preliminary analyses supported this hypothesis as light exposure was negatively related to income (r=-0.081, p=10**^-^**^5^), having at least intermediate education (r=-0.223, p=10**^-^**^42^), and population of the place of residence (r=-0.197, p=10**^-^**^33^); while both higher income and having at least intermediate education was related to better scores on all three mental wellbeing measures (absolute r=0.041-0.149, p**max**=0.004). (Population of the size of residence, however, was slightly but significantly related to both lower WHO5 [r=-0.028, p=0.02] and higher BDI [r=0.04, p=9*10**^-^**^4^] scores, and unrelated to perceived stress.) We solved this issue by controlling for various possible confounding factors, including the type of settlements respondents reside in. Overall, we failed to find evidence for an association between light exposure and mental wellbeing. Scores on the Beck Depression Inventory were unrelated to light exposure regardless of the covariates used. Both perceived stress and, paradoxically, wellbeing was slightly negatively related to light exposure, but these associations lost their significance after controls for income and education. Our findings do not support the hypothesis that light exposure is strongly, causally related to mental wellbeing. The only such association exists with perceived stress, it is weak, and it is fully accounted for by the effects of education and income.

### Social jetlag

Social jetlag, the misalignment between biological time and the timing of social obligations during working days, has been shown to contribute to the risk of a host of chronic diseases, including leading causes of death ^29^. Our cross-sectional sample was not designed to detect the effects of social jetlag on health, as these effects take a long time the develop and the social jetlag that contributed to the ailments our respondents reported may be decades in the past. We were, however, able to pinpoint both demographic and health-related factors which affect social jetlag and thus may contribute to future disease through this pathway. We found that social jetlag is lower in women and in older participants, with some evidence for a diminishing sex difference in older age. Social jetlag exhibited similar geographic associations to chronotype. Most anthropometric/biomedical variables were associated with less social jetlag; however, these associations are strongly confounded by age. After accounting for age, sex, education and income, 15 variables remained associated with social jetlag with at least nominal significance. However, the direction of associations was not consistent with the hypothesis that having health problems confers an additional risk for further illness through increasing social jetlag, as most health problems were associated with lower, not higher levels of social jetlag.

All correlations with social jetlag may partially reflect the fact that the earlier chronotypes observed in both women and older individuals are better aligned with social time. Early chronotype was a strong predictor of low social jetlag (with an estimated 0.36 hour more social jetlag for a one hour later chronotype, p=10^−113^, net of demographic predictors). Consequently, controlling chronotype drastically altered the landscape of variables associated with social jetlag. While the effects of age, childcare and income remained similar, geographical effects were almost completely eliminated, the effect of education changed signs from positive to negative, and most health associations also disappeared when controlling for chronotype. After this control, only smoking and better self-rated mental health were convincingly associated with higher levels of social jetlag.

These observations suggest that certain factors – such as age, childcare or having a higher education – are associated with less risk for social jetlag and, consequently, its deleterious effects on health. However, most of these associations are due to differences in chronotype, which is strongly predictive of the amount of social jetlag individuals experience. For individuals with identical chronotypes, only a few correlates of social jetlag were found.

### Limitations and summary

Although our study is based on a large and nationally representative sample, which provides considerable statistical power, some issues need to be considered in the interpretation of our findings. First, our estimate of chronotype relied on self-reports. Self-reports of chronotypes are known to correlate reasonably, but not perfectly objective measurements of daily rhythms ^62–64^, possibly resulting in biases in our findings. Second, as self-reported chronotype using the standard MCTQ can only be calculated for participants who report regular work schedules on working days (e.g. no permanent shift work) and the ability to freely adjust their sleeping times on free days. Participants without these characteristics are not represented in our analyses. An alternative way of assessing these types of associations might be performed by using the morningness-eveningness concept, relying on items asking the preferred time schedules of the respondents. This type of replication could further strengthen our current findings.

In sum, our findings provide evidence that chronotype and social jetlag are strongly affected by demographic and geographical factors, even if their effect is studied independently. Evidence for associations with health-related factors, however, are much more limited and mainly consist of associations with chronotype, but not chronotype-adjusted social jetlag. Importantly, contrary to some previous findings in smaller samples, we found no evidence for an association between light exposure and mental health.

## Acknowledgements

This publication has been supported by the National Research, Development and Innovation Fund of the Ministry of Innovation and Technology (grant IDs: OTKA PD 138935, TKP2021- EGA-25 and ÚNKP-22-3-II).

