## Supplementary material for "Daily rhythms, demographics and health: findings from a nationally representative survey": hungarostudy_analyze_clean.m

```
%reads Hungarostudy data and calculates demographic and health-related
%correlates


clear all
% docid='1F1TF5yXyPbtE8zi4GEpGJFXObxXkuLO1';
% sheet_name='Hungarostudy 2021 adatb�zis 20211014';
% url_name=sprintf('https://docs.google.com/spreadsheets/d/%s/gviz/tq?tqx=out:csv&sheet=%s',docid, sheet_name);
% data=webread(url_name);
% 
data=readtable('C:\Users\Ujma P�ter\OneDrive\Studies\Hungarostudy chronotype\Hungarostudy 2021 adatb�zis 20211014.csv');
settldata=importdata('C:\Users\Ujma P�ter\OneDrive\Studies\Hungarostudy chronotype\settlements.txt');
addpath('C:\Users\Ujma P�ter\OneDrive\MATLAB scripts')
%Manually fixing entries
%Mikolc
settldata.data(settldata.data==47.9804)=48.104167;
settldata.data(settldata.data==20.9354)=20.791667;

%Debrecen
settldata.data(settldata.data==47.7675)=47.52997;
settldata.data(settldata.data==21.24)=21.63916;

%Ny�regyh�za
settldata.data(settldata.data==47.95)=47.95306;
settldata.data(settldata.data==21.7167)=21.72713;
 
%Szeged
settldata.data(settldata.data==46.318)=46.255;
settldata.data(settldata.data==20.027)=20.145;

%Gy�r
settldata.data(settldata.data==47.6076)=47.6842;
settldata.data(settldata.data==17.781)=17.6344;

%P�cs
settldata.data(settldata.data==46.0967)=46.07125;
settldata.data(settldata.data==18.2068)=18.23311;


irsz=settldata.textdata(:,1);
irsz=cellfun(@str2num,irsz);
for i=1:size(data,1)
    idx=find(data.irsz_m(i)==irsz);
    if ~isempty(idx)
    idx=idx(1); %many postal codes are issued for multiple villages. Database doesn't use KSH codes so the best we can do is assume the first alphabetically. These should be close to each other.
    if data.irsz_m(i)<2000 %respondent in Budapest, let's take citywide data instead of district data for area and population stats (but not for coordinates)
    data.Latitude(i)=settldata.data(idx,1);
    data.Longitude(i)=settldata.data(idx,2);
    data.Kshcode(i)=settldata.data(idx,3);
    data.Jogallas(i)=settldata.data(idx,4);
    data.Terulet(i)=52514; 
    data.Nepesseg(i)=1723836; %Wikipedia data for 2021. 01. 01.
    data.Lakszam(i)=924664; %data: https://hu.wikipedia.org/wiki/Magyarorsz%C3%A1g_legnagyobb_telep%C3%BCl%C3%A9sei_a_lak%C3%A1sok_sz%C3%A1ma_szerint
    else 
    data.Latitude(i)=settldata.data(idx,1);
    data.Longitude(i)=settldata.data(idx,2);
    data.Kshcode(i)=settldata.data(idx,3);
    data.Jogallas(i)=settldata.data(idx,4);
    data.Terulet(i)=settldata.data(idx,5);
    data.Nepesseg(i)=settldata.data(idx,6);
    data.Lakszam(i)=settldata.data(idx,7);
    end
    end
end

% data.Joga_kat=categorical(data.Jogallas);
data.Joga_kat(data.Jogallas==1)={'County seat'};
data.Joga_kat(data.Jogallas==2)={'Budapest'};
data.Joga_kat(data.Jogallas==3)={'Town'};
data.Joga_kat(data.Jogallas==4)={'Town'};
data.Joga_kat(data.Jogallas==5)={'Large village'};
data.Joga_kat(data.Jogallas==6)={'Village'};
data.Kor=2021-data.sz__v;
data.isk_v_g=categorical(data.isk_v_g);
data.Isk3=categorical(data.Isk3);
data.nem=categorical(data.nem);

%converting chronotype to hour format
splittime=datevec(data.MSFsc,'yyyy.dd.mm HH:MM:SS');
splittime2=splittime(:,4)+splittime(:,5)/60+splittime(:,6)/3600;
splittime2(splittime2>12)=splittime2(splittime2>12)-24; %MSFsc after noon (but before midnight) is expressed as before midnight
data.MSFsc_corr=splittime2;

%creating settlement-level data
%starting at individual post codes
coordinates=unique(data.irsz_m(data.irsz_m>1999));
settltable=table;
%manually assigning Budapest
settltable.count(1)=sum(data.irsz_m<2000);  
settltable.MSFsc_corr(1)=nanmean(data.MSFsc_corr(data.irsz_m<2000));  
settltable.Longitude(1)=nanmean(data.Longitude(data.irsz_m<2000));  
settltable.Latitude(1)=nanmean(data.Latitude(data.irsz_m<2000));  
settltable.Nepesseg(1)=nanmean(data.Nepesseg(data.irsz_m<2000));  
settltable.Lakszam(1)=nanmean(data.Lakszam(data.irsz_m<2000));  
settltable.Terulet(1)=nanmean(data.Terulet(data.irsz_m<2000));
settltable.Irsz(1)=1000;
for settl=2:length(coordinates)
      settltable.count(settl)=sum(data.irsz_m==coordinates(settl)); %number of people in this postal code
      settltable.MSFsc_corr(settl)=nanmean(data.MSFsc_corr(data.irsz_m==coordinates(settl)));  
      settltable.Longitude(settl)=nanmean(data.Longitude(data.irsz_m==coordinates(settl)));  
      settltable.Latitude(settl)=nanmean(data.Latitude(data.irsz_m==coordinates(settl)));  
      settltable.Nepesseg(settl)=nanmean(data.Nepesseg(data.irsz_m==coordinates(settl)));  
      settltable.Lakszam(settl)=nanmean(data.Lakszam(data.irsz_m==coordinates(settl)));  
      settltable.Terulet(settl)=nanmean(data.Terulet(data.irsz_m==coordinates(settl)));  
      settltable.Irsz(settl)=nanmean(data.irsz_m(data.irsz_m==coordinates(settl)));
end

%fixing settltable by combining settlements with the same number of
%apartments (this is the only value that is truly unique). But first we fix
%small settlements with accidentally the same number of apartments by
%adding 1 to the values of duplicates (we don't use these data anymore
settltable = sortrows(settltable,'Lakszam','ascend');
nochange=find(diff(settltable.Lakszam)==0); %subsequent settlements have the same number of apartments
nochange(nochange>202)=[]; %by manual checking I saw that this value is the last one which is not a duplicate
settltable.Lakszam(nochange)=settltable.Lakszam(nochange)-1;

settltable2=table;
%pooling across multiple post codes per city
% [coordinates2,ia,ib]=unique([settltable.Latitude settltable.Longitude],'rows');
coordinates2=unique(settltable.Lakszam);
for settl=1:length(coordinates2)
%       idx=find(settltable.Latitude==coordinates2(settl,1) & settltable.Longitude==coordinates2(settl,2)); %indices to all cases
      idx=find(settltable.Lakszam==coordinates2(settl,1)); %indices to all cases
      count=nansum(settltable.count(idx)); %number of people at this latitude
      settltable2.count(settl)=count; %number of people at this latitude
      settltable2.MSFsc_corr(settl)=nansum(settltable.MSFsc_corr(idx).*settltable.count(idx))/count; %weighted average at this latitude
      settltable2.Longitude(settl)=nanmean(settltable.Longitude(idx)); %average at this latitude, should not change, fractional values are a sign of error
      settltable2.Latitude(settl)=nanmean(settltable.Latitude(idx)); %average at this latitude, should not change, fractional values are a sign of error
      settltable2.Nepesseg(settl)=nanmean(settltable.Nepesseg(idx)); %average at this latitude, should not change, fractional values are a sign of error
      settltable2.Lakszam(settl)=nanmean(settltable.Lakszam(idx)); %average at this latitude, should not change, fractional values are a sign of error
      settltable2.Terulet(settl)=nanmean(settltable.Terulet(idx)); %average at this latitude, should not change, fractional values are a sign of error
end
settltable2.MSFsc_corr(settltable2.MSFsc_corr==0)=NaN;
settltable2.Nepesseg=settltable2.Nepesseg/1000;


%correcting social jetlag
data.SJLrel(data.SJLrel>0.5)=data.SJLrel(data.SJLrel>0.5)-1;
data.SJLrel(data.SJLrel<-0.5)=abs(data.SJLrel(data.SJLrel<-0.5)+0.5);
data.SJLrel=data.SJLrel*24;
data.SJLabs=abs(data.SJLrel);

%deleting chronotype and SJL observations with data issues
data.MSFsc_corr(data.kizaras~=1)=NaN;

%setting up variables
% data.sex=categorical(data.nem,1:2,{'Male' 'Female'});
%coding income
incomebrackets=[15 40 60 80 100 120 140 175 225 275 325 375 425 475 550];
data.income=nan(size(data,1),1);
for br=1:length(incomebrackets)
data.income(data.j_v_nett_szem==br)=incomebrackets(br);
end
data.income=data.income/100; %expressing inhabitants in hundreds of thousands to get more interpretable estimates

%coding illnesses
illvar=find(contains(data.Properties.VariableNames,'betegs_g') & ~contains(data.Properties.VariableNames,'_E'));
illnesses={'t1db', 't2db', 'liver', 'asthma', 'other_resp', 'allergy', 'ulcer', 'other_digest', 'kidney', 'rheuma', 'other_musc', 'trafficacc', 'workacc', 'homeacc', 'highbp', 'cancer', 'psychiatric', 'anxiety', 'heart', 'cerebrovasc', 'covid', 'vacc', 'otherill'};
illnesses2={'Type 1 diabetes', 'Type 2 diabetes', 'Liver disease', 'Asthma', 'Other resp. illness', 'Allergy', 'Ulcer', 'Other digestive', 'Kidney disease', 'Rheuma', 'Other musculoskeletal', 'Traffic accident', 'Work accident', 'Home accident', 'High BP', 'Cancer', 'Psychiatric', 'Anxiety', 'Heart disease', 'Cerebrovascular', 'COVID-19', 'C19 vaccination' 'Other disease'};
illnesscat={'No','Yes','Yes','Yes'};
for ill=1:length(illvar)
   eval(['data.' illnesses{ill} '=categorical(data.betegs_g' num2str(ill) ',0:3,illnesscat);']);
end
%coding phq symptoms
phqvar=find(contains(data.Properties.VariableNames,'PHQ') & ~contains(data.Properties.VariableNames,'Sum'));
phqitems={'Stomach pain', 'Back pain', 'Limb pain', 'Headache', 'Chest pain', 'Dizzyness', 'Fainting and weakness', 'Palpitation', 'Breathing difficulty', 'Sexual problems', 'Constipation or diarrhea', 'Digestion problems', 'Tiredness', 'Sleep problems'};
%coding education
data.edu=data.Isk3;
%coding phq
phqcat={'0','0','1','1'};
for phq=1:length(phqvar)
   eval(['data.PHQc' num2str(phq) '=categorical(data.PHQ' num2str(phq) ',0:3,phqcat);']);
end
%coding cohabitation
cohabvar=find(contains(data.Properties.VariableNames,'egy_tt_l'));
% data.cohabitation=undefined(size(data,1),1);
data.cohab1=nan(size(data,1),1);
data.cohab1(sum(table2array(data(:,cohabvar)),2)>0)=1;
data.cohab1(sum(table2array(data(:,cohabvar)),2)==0)=0;
data.cohabitation=categorical(data.cohab1,0:1,{'Non-cohabiting','Cohabiting'});
data.cohab2=nan(size(data,1),1);
data.cohab2(data.egy_tt_l3>0 & data.Kor-data.sz_l_s4<6)=1;
data.cohab2(data.egy_tt_l3==0 | data.Kor-data.sz_l_s4>6)=0;
data.cohabchild=categorical(data.cohab2,0:1,{'Non-cohabiting','Cohabiting'});
%cohabitation over
data.orientation=categorical(data.sex7,1:4,{'Heterosexual' 'Homosexual' 'Bisexual' 'Asexual'});
data.ethn=categorical(data.etnikum2,0:1,{'Non-gypsy' 'Gypsy'});
data.religious=data.vall_s1+data.vall_s2;
data.physical=data.fiz_akt1+data.fiz_akt2;
data.numberdays=data.bet_nap;
data.psychotrop=categorical(data.nyugtat_,1:7,{'Rare use' 'Rare use' 'Regular use' 'Regular use' 'Regular use' 'Regular use' 'Regular use'});
data.suicide=data.suic1+data.suic2+data.suic3;
data.lightexposure=nan(size(data,1),1);
regular=data.MCQ10_1==1;
irregular=data.MCQ10_1==0;
data.lightexposure(regular)=data.MCQ10_2(regular).*nansum([data.v5(regular) data.v6(regular)/60],2)+(7-data.MCQ10_2(regular)).*nansum([data.v7(regular) data.v8(regular)/60],2);
data.lightexposure(irregular)=data.MCQ10_2(irregular).*nansum([data.v5(irregular) data.v6(irregular)/60],2)+(7-data.MCQ10_2(irregular)).*nansum([data.v7(irregular) data.v8(irregular)/60],2);
data.Nepesseg=data.Nepesseg/100000; %expressing inhabitants in hundred thousands to get more interpretable estimates
data.MSFsc_min=data.MSFsc_corr*60; %expressing chronotype as minutes relative to midnight
for id=1:length(data.K_RDEZ_SIDEJEKEZDETE)
    admintimes(:,id)=nan(6,1);
    if size(data.K_RDEZ_SIDEJEKEZDETE{id},1)>0
    admintimes(:,id)=datevec(data.K_RDEZ_SIDEJEKEZDETE(id));
    end
end
data.admintime=[admintimes(4,:)+admintimes(5,:)/60+admintimes(6,:)/3600]'; %expressing the start of administration time as fractional hours from midnight
data.admintime=data.admintime-12; %relative to noon
data.admintime_squared=data.admintime.^2;

aisvar=find(contains(data.Properties.VariableNames,'AIS8'));
data.AISSum=nansum(table2array(data(:,aisvar)),2);
%coding alcohol consumption
data.alc1=nan(size(data,1),1);
data.alc1(data.alkohol1==1)=0;
data.alc1(data.alkohol1==2)=1;
data.alc1(data.alkohol1==3)=3;
data.alc1(data.alkohol1==4)=10;
data.alc1(data.alkohol1==5)=16;
data.alc2=nan(size(data,1),1);
data.alc2(data.alkohol2==1)=1.5;
data.alc2(data.alkohol2==2)=3.5;
data.alc2(data.alkohol2==3)=5.5;
data.alc2(data.alkohol2==4)=8;
data.alc2(data.alkohol2==5)=10;
data.alcohol=nan(size(data,1),1);
data.alcohol(data.alc1==0)=0;
data.alcohol(data.alc1~=0)=data.alc1(data.alc1~=0).*data.alc2(data.alc1~=0);
%coding smoking
data.smoker=nan(size(data,1),1);
data.smoker(data.doh_ny1==1)=1;
data.smoker(data.doh_ny1==2)=1;
data.smoker(data.doh_ny1==3)=1;
data.smoker(data.doh_ny1==4)=0;
data.smoker(data.doh_ny1==5)=0;
data.smoker=categorical(data.smoker);

%Z-transforming continuous variables to get better coefficients
data.BDIz=(data.BDI-nanmean(data.BDI))/nanstd(data.BDI);
data.religiousz=(data.religious-nanmean(data.religious))/nanstd(data.religious);
data.physicalz=(data.physical-nanmean(data.physical))/nanstd(data.physical);
data.numberdaysz=(data.numberdays-nanmean(data.numberdays))/nanstd(data.numberdays);
data.PSS10Sumz=(data.PSS10Sum-nanmean(data.PSS10Sum))/nanstd(data.PSS10Sum);
data.PHQSumz=(data.PHQSum-nanmean(data.PHQSum))/nanstd(data.PHQSum);
data.WHO5z=(data.WHO5-nanmean(data.WHO5))/nanstd(data.WHO5);
data.AISSumz=(data.AISSum-nanmean(data.AISSum))/nanstd(data.AISSum);


%variables
continuous={'Kor' 'BMI' 'alcohol' 'religiousz' 'income' 'physicalz' 'numberdays' 'BDIz' 'PSS10Sumz' 'PHQSumz' 'Latitude' 'Longitude' 'Nepesseg' 'WHO5z' 'AISSumz'}; %excluded suicide, too many missing cases. 'income' is a borderline case, 2500 missing observations but a significant association if left in
categoric={'smoker' 'edu' 'nem' 'cohabitation' 'orientation' 'ethn' 'psychotrop' 'cohabchild'}; %about cohabchild: it currently compares people living with small children to everybody (those with older kids, living alone etc)
for phq=1:length(phqvar)
    categoric=[categoric ['PHQc' num2str(phq)]];
end
for ill=1:length(illnesses)
    if ~contains(illnesses{ill},'other')
    categoric=[categoric illnesses{ill}];
    end
end


%creating Wilkinson notation for the full regression model
wilkinson=[];
for v=1:length(continuous)
    wilkinson=[wilkinson '+' continuous{v}];
end
for v=1:length(categoric)
    wilkinson=[wilkinson '+' categoric{v}];
end
wilkinson(1)='~';


%Chronotype
demogr_chrono=fitlm(data,'MSFsc_min~Kor*nem+cohabchild+cohabitation+orientation+religiousz+ethn+edu');
demogr_chrono_income=fitlm(data,'MSFsc_min~Kor*nem+cohabchild+cohabitation+orientation+religiousz+ethn+edu+income');
geogr_chrono1=fitlm(data,'MSFsc_min~Longitude+Latitude+Nepesseg');
geogr_chrono2=fitlm(data,'MSFsc_min~Kor*nem+Longitude+Latitude+Nepesseg');
geogr_chrono3=fitlm(data,'MSFsc_min~Kor*nem+edu+income+Longitude+Latitude+Nepesseg');


%Social jetlag
demogr_sjl=fitlm(data,'SJLrel~Kor*nem+cohabchild+cohabitation+orientation+religiousz+ethn+edu');
demogr_sjl_income=fitlm(data,'SJLrel~Kor*nem+cohabchild+cohabitation+orientation+religiousz+ethn+edu+income');
demogr_sjl_chrono=fitlm(data,'SJLrel~Kor*nem+cohabchild+cohabitation+orientation+religiousz+ethn+edu+income+MSFsc_corr');
geogr_sjl1=fitlm(data,'SJLrel~Longitude+Latitude+Nepesseg');
geogr_sjl2=fitlm(data,'SJLrel~Kor*nem+Longitude+Latitude+Nepesseg');
geogr_sjl3=fitlm(data,'SJLrel~Kor*nem+edu+income+Longitude+Latitude+Nepesseg');
geogr_sjl4=fitlm(data,'SJLrel~Kor*nem+edu+income+Longitude+Latitude+Nepesseg+MSFsc_corr');

%assembling all health variables
healthvars={'smoker' 'psychotrop' 'alcohol' 'physicalz' 'numberdays' 'BDIz' 'PHQSumz' 'WHO5z' 'AISSumz' 'BMI' 'PSS10Sumz'};
healthvars=[healthvars illnesses];
for phq=1:length(phqvar)
    healthvars=[healthvars ['PHQc' num2str(phq)]];
end
for mdl=1:length(healthvars)
    health_chrono{mdl,1}=fitlm(data,['MSFsc_min~' healthvars{mdl}]);
    health_chrono{mdl,2}=fitlm(data,['MSFsc_min~Kor*nem+' healthvars{mdl}]);
    health_chrono{mdl,3}=fitlm(data,['MSFsc_min~Kor*nem+edu+income+' healthvars{mdl}]);
    health_chrono2{mdl,1}=fitlm(data,['SJLrel~' healthvars{mdl}]);
    health_chrono2{mdl,2}=fitlm(data,['SJLrel~Kor*nem+' healthvars{mdl}]);
    health_chrono2{mdl,3}=fitlm(data,['SJLrel~Kor*nem+edu+income+' healthvars{mdl}]);
    health_chrono2{mdl,4}=fitlm(data,['SJLrel~Kor*nem+edu+income+MSFsc_corr+' healthvars{mdl}]); %an additional model which tests for interactions with perceived stress
    health_chrono2{mdl,5}=fitlm(data,['SJLrel~Kor*nem+edu+income+' healthvars{mdl} '*PSS10Sumz']); %an additional model which tests for interactions with perceived stress
    healthcoeffs.es(mdl,1)=health_chrono{mdl,1}.Coefficients.Estimate(find(contains(health_chrono{mdl,1}.CoefficientNames,healthvars{mdl})));
    healthcoeffs.es(mdl,2)=health_chrono{mdl,2}.Coefficients.Estimate(find(contains(health_chrono{mdl,2}.CoefficientNames,healthvars{mdl})));
    healthcoeffs.es(mdl,3)=health_chrono{mdl,3}.Coefficients.Estimate(find(contains(health_chrono{mdl,3}.CoefficientNames,healthvars{mdl})));
    healthcoeffs.se(mdl,1)=health_chrono{mdl,1}.Coefficients.SE(find(contains(health_chrono{mdl,1}.CoefficientNames,healthvars{mdl})));
    healthcoeffs.se(mdl,2)=health_chrono{mdl,2}.Coefficients.SE(find(contains(health_chrono{mdl,2}.CoefficientNames,healthvars{mdl})));
    healthcoeffs.se(mdl,3)=health_chrono{mdl,3}.Coefficients.SE(find(contains(health_chrono{mdl,3}.CoefficientNames,healthvars{mdl})));
    healthcoeffs.p(mdl,1)=health_chrono{mdl,1}.Coefficients.pValue(find(contains(health_chrono{mdl,1}.CoefficientNames,healthvars{mdl})));
    healthcoeffs.p(mdl,2)=health_chrono{mdl,2}.Coefficients.pValue(find(contains(health_chrono{mdl,2}.CoefficientNames,healthvars{mdl})));
    healthcoeffs.p(mdl,3)=health_chrono{mdl,3}.Coefficients.pValue(find(contains(health_chrono{mdl,3}.CoefficientNames,healthvars{mdl})));
    healthcoeffs2.es(mdl,1)=health_chrono2{mdl,1}.Coefficients.Estimate(find(contains(health_chrono2{mdl,1}.CoefficientNames,healthvars{mdl})));
    healthcoeffs2.es(mdl,2)=health_chrono2{mdl,2}.Coefficients.Estimate(find(contains(health_chrono2{mdl,2}.CoefficientNames,healthvars{mdl})));
    healthcoeffs2.es(mdl,3)=health_chrono2{mdl,3}.Coefficients.Estimate(find(contains(health_chrono2{mdl,3}.CoefficientNames,healthvars{mdl})));
    healthcoeffs2.es(mdl,4)=health_chrono2{mdl,4}.Coefficients.Estimate(find(contains(health_chrono2{mdl,4}.CoefficientNames,healthvars{mdl})));
    healthcoeffs2.se(mdl,1)=health_chrono2{mdl,1}.Coefficients.SE(find(contains(health_chrono2{mdl,1}.CoefficientNames,healthvars{mdl})));
    healthcoeffs2.se(mdl,2)=health_chrono2{mdl,2}.Coefficients.SE(find(contains(health_chrono2{mdl,2}.CoefficientNames,healthvars{mdl})));
    healthcoeffs2.se(mdl,3)=health_chrono2{mdl,3}.Coefficients.SE(find(contains(health_chrono2{mdl,3}.CoefficientNames,healthvars{mdl})));
    healthcoeffs2.se(mdl,4)=health_chrono2{mdl,4}.Coefficients.SE(find(contains(health_chrono2{mdl,4}.CoefficientNames,healthvars{mdl})));
    healthcoeffs2.p(mdl,1)=health_chrono2{mdl,1}.Coefficients.pValue(find(contains(health_chrono2{mdl,1}.CoefficientNames,healthvars{mdl})));
    healthcoeffs2.p(mdl,2)=health_chrono2{mdl,2}.Coefficients.pValue(find(contains(health_chrono2{mdl,2}.CoefficientNames,healthvars{mdl})));
    healthcoeffs2.p(mdl,3)=health_chrono2{mdl,3}.Coefficients.pValue(find(contains(health_chrono2{mdl,3}.CoefficientNames,healthvars{mdl})));
    healthcoeffs2.p(mdl,4)=health_chrono2{mdl,4}.Coefficients.pValue(find(contains(health_chrono2{mdl,4}.CoefficientNames,healthvars{mdl})));
end


%light exposure

mentalhealth={'BDIz' 'PSS10Sumz' 'WHO5z'};
for var=1:length(mentalhealth)
    wilkinson1=[mentalhealth{var} '~lightexposure'];
    wilkinson2=[mentalhealth{var} '~lightexposure+Kor*nem'];
    wilkinson3=[mentalhealth{var} '~lightexposure+Kor*nem+edu+income'];
    wilkinson4=[mentalhealth{var} '~lightexposure+Kor*nem+edu+income+Joga_kat'];
    lexpos{var,1}=fitlm(data,wilkinson1);
    lexpos{var,2}=fitlm(data,wilkinson2);
    lexpos{var,3}=fitlm(data,wilkinson3);
    lexpos{var,4}=fitlm(data,wilkinson4);
    mentalcoeffs.es(var,1)=lexpos{var,1}.Coefficients.Estimate(find(contains(lexpos{var,1}.CoefficientNames,'lightexposure')));
    mentalcoeffs.es(var,2)=lexpos{var,2}.Coefficients.Estimate(find(contains(lexpos{var,2}.CoefficientNames,'lightexposure')));
    mentalcoeffs.es(var,3)=lexpos{var,3}.Coefficients.Estimate(find(contains(lexpos{var,3}.CoefficientNames,'lightexposure')));
    mentalcoeffs.es(var,4)=lexpos{var,4}.Coefficients.Estimate(find(contains(lexpos{var,4}.CoefficientNames,'lightexposure')));
    mentalcoeffs.se(var,1)=lexpos{var,1}.Coefficients.SE(find(contains(lexpos{var,1}.CoefficientNames,'lightexposure')));
    mentalcoeffs.se(var,2)=lexpos{var,2}.Coefficients.SE(find(contains(lexpos{var,2}.CoefficientNames,'lightexposure')));
    mentalcoeffs.se(var,3)=lexpos{var,3}.Coefficients.SE(find(contains(lexpos{var,3}.CoefficientNames,'lightexposure')));
    mentalcoeffs.se(var,4)=lexpos{var,4}.Coefficients.SE(find(contains(lexpos{var,4}.CoefficientNames,'lightexposure')));
    mentalcoeffs.p(var,1)=lexpos{var,1}.Coefficients.pValue(find(contains(lexpos{var,1}.CoefficientNames,'lightexposure')));
    mentalcoeffs.p(var,2)=lexpos{var,2}.Coefficients.pValue(find(contains(lexpos{var,2}.CoefficientNames,'lightexposure')));
    mentalcoeffs.p(var,3)=lexpos{var,3}.Coefficients.pValue(find(contains(lexpos{var,3}.CoefficientNames,'lightexposure')));
    mentalcoeffs.p(var,4)=lexpos{var,4}.Coefficients.pValue(find(contains(lexpos{var,4}.CoefficientNames,'lightexposure')));
end


%calculating social jetlag effects!
for var=1:length(healthvars)
    wilkinson=['SJLrel~' healthvars{var}];
    wilkinson2=['SJLrel~Kor*nem+' healthvars{var}];
    wilkinson3=['SJLrel~Kor*nem+edu+income+' healthvars{var}];
    sjlmdl{var,1}=fitlm(data,wilkinson);
    sjlmdl{var,2}=fitlm(data,wilkinson2);
    sjlmdl{var,3}=fitlm(data,wilkinson3);
    sjlcoefs(var,1)=sjlmdl{var,1}.Coefficients.Estimate(2);
    sjlcoefs(var,2)=sjlmdl{var,2}.Coefficients.Estimate(4);
    sjlcoefs(var,3)=sjlmdl{var,3}.Coefficients.Estimate(7);
    sjlse(var,1)=sjlmdl{var,1}.Coefficients.SE(2);
    sjlse(var,2)=sjlmdl{var,2}.Coefficients.SE(4);
    sjlse(var,3)=sjlmdl{var,3}.Coefficients.SE(7);
    sjlp(var,1)=sjlmdl{var,1}.Coefficients.pValue(2);
    sjlp(var,2)=sjlmdl{var,2}.Coefficients.pValue(4);
    sjlp(var,3)=sjlmdl{var,3}.Coefficients.pValue(7);
end

%modelling daily sleepiness 
% data.admintime(data.admintime<-5 | data.admintime>8.5)=NaN; %discarding observations before 7 AM or after 8:30 PM
sleepymdl=fitlm(data,'x_lmoss_g_pre~admintime+admintime_squared*MSFsc_min');


% %PLOTTING

% demographic model
demo_coefs={'Female','Age','Intermediate education','Advanced education','Cohabits w/anyone','Cohabits w/child','Homosexual','Bisexual','Asexual','Gypsy/Roma','Religiousness','Age*sex'};
demo_coefs2={'Female','Age','Income','Intermediate education','Advanced education','Cohabits w/anyone','Cohabits w/child','Homosexual','Bisexual','Asexual','Gypsy/Roma','Religiousness','Age*sex'};
demo_coefs3={'Female','Age','Chronotype','Income','Intermediate education','Advanced education','Cohabits w/anyone','Cohabits w/child','Homosexual','Bisexual','Asexual','Gypsy/Roma','Religiousness','Age*sex'};

figures(1)=figure('units','normalized','outerposition',[0 0 0.75 1]);
tiledlayout(2,3)
nexttile
h1=errorbar(2:length(demogr_chrono.CoefficientNames),demogr_chrono.Coefficients.Estimate(2:end),1.96*demogr_chrono.Coefficients.SE(2:end),1.96*demogr_chrono.Coefficients.SE(2:end),'o','LineWidth',4);
yline(0,'LineWidth',3)
set(gca,'XTick',2:length(demogr_chrono.CoefficientNames))
set(gca,'XTickLabels',demo_coefs)
set(gca,'FontSize',16)
xlim([1 length(demogr_chrono.CoefficientNames)+1])
set(gcf,'Color','w')
grid on
ylabel('Estimated chronotype difference (minutes)')
title('Demographic effects')
view(90,90)
nexttile
idx1=(data.nem=='1' & ~isnan(data.MSFsc_min));
idx2=(data.nem=='2' & ~isnan(data.MSFsc_min));
[f1,x1]=ksdensity(data.MSFsc_min(idx1)/60);
f2=ksdensity(data.MSFsc_min(idx2)/60,x1);
% area(x1,f1,'FaceColor','b','EdgeColor','b','FaceAlpha',0.5)
% hold on
% area(x1,f2,'FaceColor','r','EdgeColor','r','FaceAlpha',0.5)
plot(x1,f1,'b','LineWidth',3)
hold on
plot(x1,f2,'r','LineWidth',3)
grid on
legend({'Male' 'Female'})
xlabel('Chronotype (hours to midnight)')
ylabel('Relative density')
title('Sex and chronotype')
set(gca,'FontSize',16)
nexttile
data=sortrows(data,'MSFsc_min','ascend'); %this is to make polyfit work
idx1=(data.nem=='1' & ~isnan(data.MSFsc_min));
idx2=(data.nem=='2' & ~isnan(data.MSFsc_min));
scatter(data.Kor(idx1),data.MSFsc_min(idx1)/60,5,'b','MarkerEdgeAlpha',0.5,'MarkerFaceAlpha',0.5);
hold on
scatter(data.Kor(idx2),data.MSFsc_min(idx2)/60,5,'r','MarkerEdgeAlpha',0.5,'MarkerFaceAlpha',0.5);
fit1=fit(data.Kor(idx1),data.MSFsc_min(idx1)/60,'poly4','Normalize','on');
fit2=fit(data.Kor(idx2),data.MSFsc_min(idx2)/60,'poly4','Normalize','on');
f1=plot(fit1,'b');
f2=plot(fit2,'r');
set(f1,'LineWidth',3)
set(f2,'LineWidth',3)
%previous solutions at fitting that didn't work
% fit1=polyfit(data.Kor(idx1),data.MSFsc_min(idx1)/60,4);
% fit2=polyfit(data.Kor(idx2),data.MSFsc_min(idx2)/60,4);
% plot(data.Kor(idx1),polyval(fit1,data.Kor(idx1)))
% plot(polyval(fit2,data.Kor(idx2)))
% h1=lsline;
% h1(1,1).LineWidth=3;
% h1(1,2).LineWidth=3;
% h1(1,1).Color='r'; %lsline doesn't keep the order of the original scatterplots so this is how it's correct!
% h1(1,2).Color='b';
legend({'Male' 'Female'})
xlabel('Age')
ylabel('Chronotype (hours to midnight)')
title('Age and chronotype')
set(gca,'FontSize',16)
grid on
nexttile
idx1=(data.edu=='1' & ~isnan(data.MSFsc_min));
idx2=(data.edu=='2' & ~isnan(data.MSFsc_min));
idx3=(data.edu=='3' & ~isnan(data.MSFsc_min));
[f1,x1]=ksdensity(data.MSFsc_min(idx1)/60);
f2=ksdensity(data.MSFsc_min(idx2)/60,x1);
f3=ksdensity(data.MSFsc_min(idx3)/60,x1);
plot(x1,f1,'k','LineWidth',3)
hold on
plot(x1,f2,'r','LineWidth',3)
plot(x1,f3,'g','LineWidth',3)
grid on
legend({'Basic education' 'Intermediate education' 'Advanced education'})
xlabel('Chronotype (hours to midnight)')
ylabel('Relative density')
title('Education and chronotype')
set(gca,'FontSize',16)
xlim([-2 12])
nexttile
idx1=(data.cohabchild=='Non-cohabiting' & ~isnan(data.MSFsc_min));
idx2=(data.cohabchild=='Cohabiting' & ~isnan(data.MSFsc_min));
[f1,x1]=ksdensity(data.MSFsc_min(idx1)/60);
f2=ksdensity(data.MSFsc_min(idx2)/60,x1);
plot(x1,f1,'c','LineWidth',3)
hold on
plot(x1,f2,'m','LineWidth',3)
grid on
legend({ 'No small child in household' 'Small child in household'})
xlabel('Chronotype (hours to midnight)')
ylabel('Relative density')
title('Cohabiting with children')
set(gca,'FontSize',16)
xlim([-2 12])
nexttile
cat=tabulate(data.Joga_kat);
for c=1:size(cat,1)
    idx=(strcmp(data.Joga_kat,cat{c,1}) & ~isnan(data.MSFsc_min));
    f=ksdensity(data.MSFsc_min(idx)/60,x1);
    plot(x1,f,'LineWidth',3)
    hold on
end
legend(cat{:,1})
grid on
xlabel('Chronotype (hours to midnight)')
ylabel('Relative density')
title('Settlement types')
set(gca,'FontSize',16)
xlim([-2 12])
export_fig('Demographic effects','-jpg','-r600')
close(gcf)

% demographic model with income
figures(2)=figure('units','normalized','outerposition',[0 0 0.4 0.95]);
h1=errorbar(2:length(demogr_chrono_income.CoefficientNames),demogr_chrono_income.Coefficients.Estimate(2:end),1.96*demogr_chrono_income.Coefficients.SE(2:end),1.96*demogr_chrono_income.Coefficients.SE(2:end),'o','LineWidth',4);
yline(0,'LineWidth',3)
set(gca,'XTick',2:length(demogr_chrono_income.CoefficientNames))
set(gca,'XTickLabels',demo_coefs2)
set(gca,'FontSize',16)
xlim([1 length(demogr_chrono_income.CoefficientNames)+1])
set(gcf,'Color','w')
ylabel('Estimated chronotype difference (minutes)')
title('Demographic effects')
view(90,90)
export_fig('Demographic effects with income','-jpg','-r600')
close(gcf)


%constructing supplementary table
demo_table=cell(1,4);
coefnames={'', 'B', 'SE', 'p'};
demo_table(:,1:4)=coefnames;
demo_table(2:length(demo_coefs)+1,1)=demo_coefs;
demo_table(2:length(demo_coefs)+1,2)=num2cell(demogr_chrono.Coefficients.Estimate(2:end));
demo_table(2:length(demo_coefs)+1,3)=num2cell(demogr_chrono.Coefficients.SE(2:end));
demo_table(2:length(demo_coefs)+1,4)=num2cell(demogr_chrono.Coefficients.pValue(2:end));
demo_table(size(demo_table,1)+1,1)={'R^2'};
demo_table(size(demo_table,1),2)={demogr_chrono.Rsquared.Adjusted};
demo_table(size(demo_table,1)+1,1)={'N'};
demo_table(size(demo_table,1),2)={demogr_chrono.NumObservations};
xlswrite('Hungarostudy chronotype supplements.xlsx',demo_table,'Demographic','A1')

demo_table2=cell(1,4);
coefnames={'', 'B', 'SE', 'p'};
demo_table2(:,1:4)=coefnames;
demo_table2(2:length(demo_coefs2)+1,1)=demo_coefs2;
demo_table2(2:length(demo_coefs2)+1,2)=num2cell(demogr_chrono_income.Coefficients.Estimate(2:end));
demo_table2(2:length(demo_coefs2)+1,3)=num2cell(demogr_chrono_income.Coefficients.SE(2:end));
demo_table2(2:length(demo_coefs2)+1,4)=num2cell(demogr_chrono_income.Coefficients.pValue(2:end));
demo_table2(size(demo_table2,1)+1,1)={'R^2'};
demo_table2(size(demo_table2,1),2)={demogr_chrono_income.Rsquared.Adjusted};
demo_table2(size(demo_table2,1)+1,1)={'N'};
demo_table2(size(demo_table2,1),2)={demogr_chrono_income.NumObservations};
xlswrite('Hungarostudy chronotype supplements.xlsx',demo_table2,'Demographic with income','A1')

demo_table_sjl=cell(1,4);
coefnames={'', 'B', 'SE', 'p'};
demo_table_sjl(:,1:4)=coefnames;
demo_table_sjl(2:length(demo_coefs)+1,1)=demo_coefs;
demo_table_sjl(2:length(demo_coefs)+1,2)=num2cell(demogr_sjl.Coefficients.Estimate(2:end));
demo_table_sjl(2:length(demo_coefs)+1,3)=num2cell(demogr_sjl.Coefficients.SE(2:end));
demo_table_sjl(2:length(demo_coefs)+1,4)=num2cell(demogr_sjl.Coefficients.pValue(2:end));
demo_table_sjl(size(demo_table_sjl,1)+1,1)={'R^2'};
demo_table_sjl(size(demo_table_sjl,1),2)={demogr_sjl.Rsquared.Adjusted};
demo_table_sjl(size(demo_table_sjl,1)+1,1)={'N'};
demo_table_sjl(size(demo_table_sjl,1),2)={demogr_sjl.NumObservations};
xlswrite('Hungarostudy chronotype supplements.xlsx',demo_table_sjl,'Demographic, SJL','A1')

demo_table2_sjl=cell(1,4);
coefnames={'', 'B', 'SE', 'p'};
demo_table2_sjl(:,1:4)=coefnames;
demo_table2_sjl(2:length(demo_coefs2)+1,1)=demo_coefs2;
demo_table2_sjl(2:length(demo_coefs2)+1,2)=num2cell(demogr_sjl_income.Coefficients.Estimate(2:end));
demo_table2_sjl(2:length(demo_coefs2)+1,3)=num2cell(demogr_sjl_income.Coefficients.SE(2:end));
demo_table2_sjl(2:length(demo_coefs2)+1,4)=num2cell(demogr_sjl_income.Coefficients.pValue(2:end));
demo_table2_sjl(size(demo_table2_sjl,1)+1,1)={'R^2'};
demo_table2_sjl(size(demo_table2_sjl,1),2)={demogr_sjl_income.Rsquared.Adjusted};
demo_table2_sjl(size(demo_table2_sjl,1)+1,1)={'N'};
demo_table2_sjl(size(demo_table2_sjl,1),2)={demogr_sjl_income.NumObservations};
xlswrite('Hungarostudy chronotype supplements.xlsx',demo_table2_sjl,'Demographic with income, SJL','A1')

demo_table3_sjl=cell(1,4);
coefnames={'', 'B', 'SE', 'p'};
demo_table3_sjl(:,1:4)=coefnames;
demo_table3_sjl(2:length(demo_coefs3)+1,1)=demo_coefs3;
demo_table3_sjl(2:length(demo_coefs3)+1,2)=num2cell(demogr_sjl_chrono.Coefficients.Estimate(2:end));
demo_table3_sjl(2:length(demo_coefs3)+1,3)=num2cell(demogr_sjl_chrono.Coefficients.SE(2:end));
demo_table3_sjl(2:length(demo_coefs3)+1,4)=num2cell(demogr_sjl_chrono.Coefficients.pValue(2:end));
demo_table3_sjl(size(demo_table3_sjl,1)+1,1)={'R^2'};
demo_table3_sjl(size(demo_table3_sjl,1),2)={demogr_sjl_chrono.Rsquared.Adjusted};
demo_table3_sjl(size(demo_table3_sjl,1)+1,1)={'N'};
demo_table3_sjl(size(demo_table3_sjl,1),2)={demogr_sjl_chrono.NumObservations};
xlswrite('Hungarostudy chronotype supplements.xlsx',demo_table3_sjl,'Demogr w inc chronotype, SJL','A1')

%geographical model
geo_coefs1={'Latitude' 'Longitude' 'Population'};
geo_coefs2={'Female' 'Age' 'Latitude' 'Longitude' 'Population' 'Age*sex'};
geo_coefs3={'Female' 'Age' 'Latitude' 'Longitude' 'Population' 'Income' 'Intermediate education' 'Advanced education' 'Age*sex'};
geo_coefs4={'Female' 'Age' 'Latitude' 'Longitude' 'Population' 'Chronotype' 'Income' 'Intermediate education' 'Advanced education' 'Age*sex'};


figures(3)=figure('units','normalized','outerposition',[0 0 0.75 1]);
tiledlayout(2,2)
nexttile
h1=errorbar(2:length(geogr_chrono1.CoefficientNames),geogr_chrono1.Coefficients.Estimate(2:end),1.96*geogr_chrono1.Coefficients.SE(2:end),1.96*geogr_chrono1.Coefficients.SE(2:end),'o','LineWidth',4);
hold on
h2=errorbar(2.25:length(geogr_chrono1.CoefficientNames)+0.25,geogr_chrono2.Coefficients.Estimate(4:6),1.96*geogr_chrono2.Coefficients.SE(4:6),1.96*geogr_chrono2.Coefficients.SE(4:6),'o','LineWidth',4);
h3=errorbar(2.5:length(geogr_chrono1.CoefficientNames)+0.5,geogr_chrono3.Coefficients.Estimate(4:6),1.96*geogr_chrono3.Coefficients.SE(4:6),1.96*geogr_chrono3.Coefficients.SE(4:6),'o','LineWidth',4);
yline(0,'LineWidth',3)
xlim([1 length(geo_coefs1)+2])
set(gca,'XTick',2:length(geo_coefs1)+1)
set(gca,'XTickLabels',geo_coefs1)
set(gca,'FontSize',16)
ylabel('Estimated chronotype difference (minutes)')
title('Model coefficients')
legend({'Unadjusted' 'Sex and age adjusted' 'Income and education adjusted'},'Location','best')
set(gcf,'Color','w')
view(90,90)
nexttile
gx=geoscatter(settltable2.Latitude,settltable2.Longitude,settltable2.Nepesseg,settltable2.MSFsc_corr,'LineWidth',1);
ax=colorbar;
colormap(jet)
ylabel(ax,'Mean chronotype of settlement')
title('Map of respondents')
set(gca,'FontSize',16)
nexttile
scatter(data.Longitude,data.MSFsc_min/60,2,'k')
h1=lsline;
h1.LineWidth=4;
h1.Color='r';
xlabel('Longitude (degrees)')
ylabel('Chronotype (hours to midnight)')
title('Longitude and chronotype')
set(gca,'FontSize',16)
nexttile
scatter(data.MSFsc_min/60,data.Latitude,2,'k')
h1=lsline;
h1.LineWidth=4;
h1.Color='r';
xlabel('Chronotype (hours to midnight)')
ylabel('Latitude (degrees)')
title('Latitude and chronotype')
set(gca,'FontSize',16)
export_fig('Geographical effects','-jpg','-r600')
close(gcf)

%settlement-level WLS regression on chronotype
settltable2.Pop100k=settltable2.Nepesseg/10;
settlchrono=fitlm(settltable2,'MSFsc_corr~Latitude+Longitude+Pop100k','Weights','Pop100k');
%figure
figures(4)=figure('units','normalized','outerposition',[0 0 0.3 0.5]);
scatter(settltable2.Longitude,settltable2.MSFsc_corr,settltable2.Pop100k*30,'b','filled','MarkerFaceAlpha',0.7)
h=lsline;
h.LineWidth=4;
h.Color='b';
ylabel('Chronotype (hours to midnight)')
xlabel('Longitude (degrees)')
title('Settlement-level chronotype and longitude')
set(gca,'FontSize',16)
set(gcf,'Color','w')
legendnum=5; %number of desired size legend entries
% legenddata=sort(settltable2.Pop100k);
legenddata2=[0.5 1 5 10]*30'; %A plausible range of settlement sizes%legenddata(1:length(legenddata)/legendnum:end);
hold on
for i=1:length(legenddata2)
% ld(i)=plot(nan,nan,'o','MarkerEdgeColor','b','MarkerSize',legenddata2(i)*30); %runs but the circle sizes on the legend are incorrect
ld(i)=scatter(nan,nan,legenddata2(i),'b','filled','MarkerFaceAlpha',0.7);
hold on
end
%Legend - I can't get it right...
% [~,objh]=legend(ld,num2str(legenddata2*1000));
% objhl=findobj(objh,'type','patch');
% for i=1:length(objhl)
%     objhl(i).MarkerSize=legenddata2(i);
% end

%set legend
% [l, hobj, hout,mout]=legend(ld,num2str(legenddata2*1000));
export_fig('Settlement-level effects','-jpg','-r600')
close(gcf)


%constructing supplementary table
geo_table=cell(2,10);
coefnames={'B', 'SE', 'p'};
geo_table(2,2:10)=repmat(coefnames,1,3);
geo_table(3:length(geo_coefs1)+2,1)=geo_coefs1;
geo_table(3:length(geo_coefs1)+2,2)=num2cell(geogr_chrono1.Coefficients.Estimate(2:end));
geo_table(3:length(geo_coefs1)+2,3)=num2cell(geogr_chrono1.Coefficients.SE(2:end));
geo_table(3:length(geo_coefs1)+2,4)=num2cell(geogr_chrono1.Coefficients.pValue(2:end));
geo_table(3:length(geo_coefs1)+2,5)=num2cell(geogr_chrono2.Coefficients.Estimate(4:6));
geo_table(3:length(geo_coefs1)+2,6)=num2cell(geogr_chrono2.Coefficients.SE(4:6));
geo_table(3:length(geo_coefs1)+2,7)=num2cell(geogr_chrono2.Coefficients.pValue(4:6));
geo_table(3:length(geo_coefs1)+2,8)=num2cell(geogr_chrono3.Coefficients.Estimate(4:6));
geo_table(3:length(geo_coefs1)+2,9)=num2cell(geogr_chrono3.Coefficients.SE(4:6));
geo_table(3:length(geo_coefs1)+2,10)=num2cell(geogr_chrono3.Coefficients.pValue(4:6));
geo_table(size(geo_table,1)+1,1)={'R^2'};
geo_table(size(geo_table,1),2)={geogr_chrono1.Rsquared.Adjusted};
geo_table(size(geo_table,1)+1,1)={'N'};
geo_table(size(geo_table,1),2)={geogr_chrono1.NumObservations};
geo_table(size(geo_table,1)-1,5)={geogr_chrono2.Rsquared.Adjusted};
geo_table(size(geo_table,1),5)={geogr_chrono2.NumObservations};
geo_table(size(geo_table,1)-1,8)={geogr_chrono3.Rsquared.Adjusted};
geo_table(size(geo_table,1),8)={geogr_chrono3.NumObservations};
geo_table(1,2)={'Unadjusted'};
geo_table(1,5)={'Age and sex adjusted'};
geo_table(1,8)={'Income and education adjusted'};
xlswrite('Hungarostudy chronotype supplements.xlsx',geo_table,'Geographic','A1')

%constructing supplementary table
geo_table_sjl=cell(2,13);
coefnames={'B', 'SE', 'p'};
geo_table_sjl(2,2:13)=repmat(coefnames,1,4);
geo_table_sjl(3:length(geo_coefs1)+2,1)=geo_coefs1;
geo_table_sjl(3:length(geo_coefs1)+2,2)=num2cell(geogr_sjl1.Coefficients.Estimate(2:end));
geo_table_sjl(3:length(geo_coefs1)+2,3)=num2cell(geogr_sjl1.Coefficients.SE(2:end));
geo_table_sjl(3:length(geo_coefs1)+2,4)=num2cell(geogr_sjl1.Coefficients.pValue(2:end));
geo_table_sjl(3:length(geo_coefs1)+2,5)=num2cell(geogr_sjl2.Coefficients.Estimate(4:6));
geo_table_sjl(3:length(geo_coefs1)+2,6)=num2cell(geogr_sjl2.Coefficients.SE(4:6));
geo_table_sjl(3:length(geo_coefs1)+2,7)=num2cell(geogr_sjl2.Coefficients.pValue(4:6));
geo_table_sjl(3:length(geo_coefs1)+2,8)=num2cell(geogr_sjl3.Coefficients.Estimate(4:6));
geo_table_sjl(3:length(geo_coefs1)+2,9)=num2cell(geogr_sjl3.Coefficients.SE(4:6));
geo_table_sjl(3:length(geo_coefs1)+2,10)=num2cell(geogr_sjl3.Coefficients.pValue(4:6));
geo_table_sjl(3:length(geo_coefs1)+2,11)=num2cell(geogr_sjl4.Coefficients.Estimate(4:6));
geo_table_sjl(3:length(geo_coefs1)+2,12)=num2cell(geogr_sjl4.Coefficients.SE(4:6));
geo_table_sjl(3:length(geo_coefs1)+2,13)=num2cell(geogr_sjl4.Coefficients.pValue(4:6));
geo_table_sjl(size(geo_table_sjl,1)+1,1)={'R^2'};
geo_table_sjl(size(geo_table_sjl,1),2)={geogr_sjl1.Rsquared.Adjusted};
geo_table_sjl(size(geo_table_sjl,1)+1,1)={'N'};
geo_table_sjl(size(geo_table_sjl,1),2)={geogr_sjl1.NumObservations};
geo_table_sjl(size(geo_table_sjl,1)-1,5)={geogr_sjl2.Rsquared.Adjusted};
geo_table_sjl(size(geo_table_sjl,1),5)={geogr_sjl2.NumObservations};
geo_table_sjl(size(geo_table_sjl,1)-1,8)={geogr_sjl3.Rsquared.Adjusted};
geo_table_sjl(size(geo_table_sjl,1),8)={geogr_sjl3.NumObservations};
geo_table_sjl(1,2)={'Unadjusted'};
geo_table_sjl(1,5)={'Age and sex adjusted'};
geo_table_sjl(1,8)={'Income and education adjusted'};
geo_table_sjl(1,11)={'Income, education and chronotype adjusted'};
xlswrite('Hungarostudy chronotype supplements.xlsx',geo_table_sjl,'Geographic, SJL','A1')

%biomedical models
healthvars2={'Smoking' 'Psychotropic medication' 'Alcohol/month' 'Physical activity' 'Days of illness' 'Beck depression' 'PHQ total' 'WHO QoL' 'Athens insomnia' 'BMI' 'PSS10'};
healthvars2=[healthvars2 illnesses2 phqitems];
figures(5)=figure('units','normalized','outerposition',[0 0 0.4 0.95]);
h1=errorbar(1:length(healthcoeffs.es),healthcoeffs.es(:,1),1.96*healthcoeffs.se(:,1),1.96*healthcoeffs.se(:,1),'o','LineWidth',2);
hold on
h2=errorbar(1.25:length(healthcoeffs.es)+0.25,healthcoeffs.es(:,2),1.96*healthcoeffs.se(:,2),1.96*healthcoeffs.se(:,2),'o','LineWidth',2);
h3=errorbar(1.5:length(healthcoeffs.es)+0.5,healthcoeffs.es(:,3),1.96*healthcoeffs.se(:,3),1.96*healthcoeffs.se(:,3),'o','LineWidth',2);
yline(0,'LineWidth',3)
set(gca,'XTick',1:length(healthvars))
set(gca,'XTickLabels',healthvars2)
xlim([0 length(healthvars)+1])
set(gcf,'Color','w')
ylabel('Estimated chronotype difference (minutes)')
legend({'Unadjusted' 'Sex and age adjusted' 'Income and education adjusted'},'Location','best')
title('Anthropometric/biomedical effects')
grid on
view(90,90)
export_fig('Biomedical effects','-jpg','-r600')
close(gcf)

%constructing supplementary table
health_table=cell(2,10);
coefnames={'B', 'SE', 'p'};
health_table(2,2:10)=repmat(coefnames,1,3);
health_table(1,2)={'Unadjusted'};
health_table(1,5)={'Age and sex adjusted'};
health_table(1,8)={'Income and education adjusted'};
health_table(3:length(healthvars2)+2,1)=healthvars2;
health_table(3:length(healthvars2)+2,2)=num2cell(healthcoeffs.es(:,1));
health_table(3:length(healthvars2)+2,3)=num2cell(healthcoeffs.se(:,1));
health_table(3:length(healthvars2)+2,4)=num2cell(healthcoeffs.p(:,1));
health_table(3:length(healthvars2)+2,5)=num2cell(healthcoeffs.es(:,2));
health_table(3:length(healthvars2)+2,6)=num2cell(healthcoeffs.se(:,2));
health_table(3:length(healthvars2)+2,7)=num2cell(healthcoeffs.p(:,2));
health_table(3:length(healthvars2)+2,8)=num2cell(healthcoeffs.es(:,3));
health_table(3:length(healthvars2)+2,9)=num2cell(healthcoeffs.se(:,3));
health_table(3:length(healthvars2)+2,10)=num2cell(healthcoeffs.p(:,3));
xlswrite('Hungarostudy chronotype supplements.xlsx',health_table,'Anthropometric and biomedical','A1')

%social jetlag effects
figures(6)=figure('units','normalized','outerposition',[0 0 0.4 0.95]);
h1=errorbar(1:length(healthcoeffs2.es),healthcoeffs2.es(:,1),1.96*healthcoeffs2.se(:,1),1.96*healthcoeffs2.se(:,1),'o','LineWidth',2);
hold on
h2=errorbar(1.25:length(healthcoeffs2.es)+0.25,healthcoeffs2.es(:,2),1.96*healthcoeffs2.se(:,2),1.96*healthcoeffs2.se(:,2),'o','LineWidth',2);
h3=errorbar(1.5:length(healthcoeffs2.es)+0.5,healthcoeffs2.es(:,3),1.96*healthcoeffs2.se(:,3),1.96*healthcoeffs2.se(:,3),'o','LineWidth',2);
h4=errorbar(1.75:length(healthcoeffs2.es)+0.75,healthcoeffs2.es(:,4),1.96*healthcoeffs2.se(:,4),1.96*healthcoeffs2.se(:,4),'o','LineWidth',2);
yline(0,'LineWidth',3)
set(gca,'XTick',1:length(healthvars))
set(gca,'XTickLabels',healthvars2)
xlim([0 length(healthvars)+1])
set(gcf,'Color','w')
ylabel('Estimated social jetlag difference (hours)')
title('Social jetlag effects')
grid on
view(90,90)
legend({'Unadjusted' 'Sex and age adjusted' 'Income and education adjusted' 'Income, education and chronotype adjusted'},'Location','best')
export_fig('Biomedical effects on social jetlag','-jpg','-r600')
close(gcf)

%constructing supplementary table
sjl_table=cell(2,13);
coefnames={'B', 'SE', 'p'};
sjl_table(2,2:13)=repmat(coefnames,1,4);
sjl_table(1,2)={'Unadjusted'};
sjl_table(1,5)={'Age and sex adjusted'};
sjl_table(1,8)={'Income and education adjusted'};
sjl_table(1,11)={'Income,  education and chronotype adjusted'};
sjl_table(3:length(healthvars2)+2,1)=healthvars2;
sjl_table(3:length(healthvars2)+2,2)=num2cell(healthcoeffs2.es(:,1));
sjl_table(3:length(healthvars2)+2,3)=num2cell(healthcoeffs2.se(:,1));
sjl_table(3:length(healthvars2)+2,4)=num2cell(healthcoeffs2.p(:,1));
sjl_table(3:length(healthvars2)+2,5)=num2cell(healthcoeffs2.es(:,2));
sjl_table(3:length(healthvars2)+2,6)=num2cell(healthcoeffs2.se(:,2));
sjl_table(3:length(healthvars2)+2,7)=num2cell(healthcoeffs2.p(:,2));
sjl_table(3:length(healthvars2)+2,8)=num2cell(healthcoeffs2.es(:,3));
sjl_table(3:length(healthvars2)+2,9)=num2cell(healthcoeffs2.se(:,3));
sjl_table(3:length(healthvars2)+2,10)=num2cell(healthcoeffs2.p(:,3));
sjl_table(3:length(healthvars2)+2,11)=num2cell(healthcoeffs2.es(:,4));
sjl_table(3:length(healthvars2)+2,12)=num2cell(healthcoeffs2.se(:,4));
sjl_table(3:length(healthvars2)+2,13)=num2cell(healthcoeffs2.p(:,4));
xlswrite('Hungarostudy chronotype supplements.xlsx',sjl_table,'Anthropom. and biomedical, SJL','A1')

%light exposure effects
mentalhealth2={'Depression (BDI)' 'Stress (PSS10)' 'Wellbeing (WHO5)'};
figures(7)=figure('units','normalized','outerposition',[0 0 0.4 0.95]);
tiledlayout(3,1)
for var=1:length(mentalhealth)
nexttile
h1=errorbar(1:length(mentalcoeffs),mentalcoeffs.es(var,1),1.96*mentalcoeffs.se(var,1),1.96*mentalcoeffs.se(var,1),'o','LineWidth',2);
hold on
h2=errorbar(1.2:length(mentalcoeffs)+0.2,mentalcoeffs.es(var,2),1.96*mentalcoeffs.se(var,2),1.96*mentalcoeffs.se(var,2),'o','LineWidth',2);
h3=errorbar(1.4:length(mentalcoeffs)+0.4,mentalcoeffs.es(var,3),1.96*mentalcoeffs.se(var,3),1.96*mentalcoeffs.se(var,3),'o','LineWidth',2);
h4=errorbar(1.6:length(mentalcoeffs)+0.6,mentalcoeffs.es(var,4),1.96*mentalcoeffs.se(var,4),1.96*mentalcoeffs.se(var,4),'o','LineWidth',2);
yline(0,'LineWidth',3)
% set(gca,'XTick',1:length(healthvars))
% set(gca,'XTickLabels',healthvars2)
ylim([-0.01 0.01])
set(gcf,'Color','w')
ylabel('Estimated score difference (SD/h)')
legend({'Unadjusted' 'Sex and age adjusted' 'Income and education adjusted' 'Residence adjusted'},'Location','best')
title(mentalhealth2{var})
grid on
view(90,90)
end
export_fig('Light exposure and mental health','-jpg','-r600')
close(gcf)

%constructing supplementary table
lexpo_table=cell(2,13);
coefnames={'B', 'SE', 'p'};
lexpo_table(2,2:13)=repmat(coefnames,1,4);
lexpo_table(1,2)={'Unadjusted'};
lexpo_table(1,5)={'Age and sex adjusted'};
lexpo_table(1,8)={'Income and education adjusted'};
lexpo_table(1,11)={'Residence adjusted'};
lexpo_table(3:length(mentalhealth2)+2,1)=mentalhealth2;
lexpo_table(3:length(mentalhealth2)+2,2)=num2cell(mentalcoeffs.es(:,1));
lexpo_table(3:length(mentalhealth2)+2,3)=num2cell(mentalcoeffs.se(:,1));
lexpo_table(3:length(mentalhealth2)+2,4)=num2cell(mentalcoeffs.p(:,1));
lexpo_table(3:length(mentalhealth2)+2,5)=num2cell(mentalcoeffs.es(:,2));
lexpo_table(3:length(mentalhealth2)+2,6)=num2cell(mentalcoeffs.se(:,2));
lexpo_table(3:length(mentalhealth2)+2,7)=num2cell(mentalcoeffs.p(:,2));
lexpo_table(3:length(mentalhealth2)+2,8)=num2cell(mentalcoeffs.es(:,3));
lexpo_table(3:length(mentalhealth2)+2,9)=num2cell(mentalcoeffs.se(:,3));
lexpo_table(3:length(mentalhealth2)+2,10)=num2cell(mentalcoeffs.p(:,3));
lexpo_table(3:length(mentalhealth2)+2,11)=num2cell(mentalcoeffs.es(:,4));
lexpo_table(3:length(mentalhealth2)+2,12)=num2cell(mentalcoeffs.se(:,4));
lexpo_table(3:length(mentalhealth2)+2,13)=num2cell(mentalcoeffs.p(:,4));
xlswrite('Hungarostudy chronotype supplements.xlsx',lexpo_table,'Light exposure','A1')


%descriptive statistics
allvars={'Kor' 'nem' 'cohabchild' 'cohabitation' 'orientation' 'religiousz' 'ethn' 'edu' 'Longitude' 'Latitude' 'Nepesseg' 'income'};
allvars2={'Age' 'Sex' 'Cohabits w/child' 'Cohabits w/anyone' 'Sexual orientation' 'Religiousness' 'Ethnicity' 'Education' 'Longitude' 'Latitude' 'Population' 'Income'};
allvars=[allvars healthvars];
allvars2=[allvars2 healthvars2];
descr_table1={};
descr_table1{1,2}='Category';
descr_table1{1,3}='Count';
descr_table1{1,4}='%';
descr_table1{1,5}='Valid N';
descr_table2={};
descr_table2{1,2}='Valid N';
descr_table2{1,3}='Mean';
descr_table2{1,4}='SD';
descr_table2{1,5}='Minimum';
descr_table2{1,6}='Maximum';
row1=2;
row2=2;
for var=1:length(allvars)
    descr={};
    if strcmp(class(eval(['data.' allvars{var}])),'categorical')
    descr_table1{row1,1}=allvars2{var};
    descr=tabulate(eval(['data.' allvars{var}]));
    descr_table1(row1:row1+size(descr,1)-1,2:4)=descr;
    descr_table1{row1+size(descr,1)-1,5}=sum(cell2mat(descr(:,2)));
    row1=row1+size(descr,1);
    elseif strcmp(class(eval(['data.' allvars{var}])),'double')
    descr_table2{row2,1}=allvars2{var};
    statdat=eval(['data.' allvars{var}]);
    descr{1,1}=sum(~isnan(statdat));
    descr{1,2}=nanmean(statdat);
    descr{1,3}=nanstd(statdat);
    descr{1,4}=min(statdat);
    descr{1,5}=max(statdat);
    descr_table2(row2:row2+size(descr,1)-1,2:6)=descr;
    row2=row2+size(descr,1);
    end
end
xlswrite('Hungarostudy chronotype descriptive statistics.xlsx',descr_table1,'Categorical variables')
xlswrite('Hungarostudy chronotype descriptive statistics.xlsx',descr_table2,'Continuous variables')

% %smoothed admin time vs. tiredness vs. chronotype
data2=sortrows(data,"admintime");
idx1=(~isnan(data2.admintime) & ~isnan(data2.x_lmoss_g_pre) & data2.MSFsc_min<nanmean(data2.MSFsc_min));
idx2=(~isnan(data2.admintime) & ~isnan(data2.x_lmoss_g_pre) & data2.MSFsc_min>nanmean(data2.MSFsc_min));
sleepyfit1=fit(data2.admintime(idx1),data2.x_lmoss_g_pre(idx1),'poly4');
sleepyfit2=fit(data2.admintime(idx2),data2.x_lmoss_g_pre(idx2),'poly4');
% sleepyfit1b=fit(data2.admintime(idx1),data2.x_lmoss_g_pre(idx1),'poly2');
% sleepyfit2b=fit(data2.admintime(idx2),data2.x_lmoss_g_pre(idx2),'poly2');
figures(8)=figure('units','normalized','outerposition',[0 0 0.5 0.8]);
h1=plot(sleepyfit1,'b');
% h1b=plot(sleepyfit1b,'b','FaceAlpha',0.5);
set(h1,'LineWidth',5)
hold on
h2=plot(sleepyfit2,'r');
% h2b=plot(sleepyfit2b,'b','FaceAlpha',0.5);
set(h2,'LineWidth',5)
xlim([-5 8.5])
xlabel('Administration time relative to noon (h)')
ylabel('Self-reported sleepiness')
grid on
legend({'Earlier than average chronotype' 'Later than average chronotype'},'Location','southeast')
dim=[.1 0 .1 .1];
title('Daily course of sleepiness')
set(gca,'FontSize',16)
set(gcf,'Color','w')
export_fig('Daily course of sleepiness vs. chronotype','-jpg','-r600')
close(gcf)

%zero-order correlations of light exposure, SES and wellbeing
data.interm_edu=nan(size(data,1),1);
data.interm_edu(data.edu=='1')=0;
data.interm_edu(data.edu=='2')=1;
data.interm_edu(data.edu=='3')=1;

[r,p]=corr(data.lightexposure,data.income,'rows','complete');
[r,p]=corr(data.lightexposure,data.interm_edu,'rows','complete');
[r,p]=corr(data.lightexposure,data.Nepesseg,'rows','complete');

[r,p]=corr(data.PSS10Sumz,data.income,'rows','complete');
[r,p]=corr(data.PSS10Sumz,data.interm_edu,'rows','complete');
[r,p]=corr(data.PSS10Sumz,data.Nepesseg,'rows','complete');

[r,p]=corr(data.WHO5z,data.income,'rows','complete');
[r,p]=corr(data.WHO5z,data.interm_edu,'rows','complete');
[r,p]=corr(data.WHO5z,data.Nepesseg,'rows','complete');

[r,p]=corr(data.BDIz,data.income,'rows','complete');
[r,p]=corr(data.BDIz,data.interm_edu,'rows','complete');
[r,p]=corr(data.BDIz,data.Nepesseg,'rows','complete');
```
